# ^1^H *R*_1ρ_ Relaxation Identifies a Hidden Intermediate in DNA Base-Pairing

**DOI:** 10.1101/2025.10.03.677141

**Authors:** Rubin Dasgupta, Christian Steinmetzger, Julian Ilgen, Katja Petzold

**Author notes:** Correspondence to Prof. Katja Petzold. Equal contribution (Rubin Dasgupta, Christian Steinmetzger). Institute of Organic Chemistry, University of Regensburg, Universitätsstraße 31, 93053 Regensburg, Germany (Julian Ilgen).

## Abstract

^1^H *R*_1ρ_ Relaxation dispersion (RD) NMR experiments provide valuable atomic-level insights into transient, high-energy conformational states of biomolecules. However, cross-relaxation artifacts can hamper its interpretation and therefore limiting broader adoption. This study explicitly quantifies cross-relaxation effects on ^1^H *R*_1*ρ*_ relaxation rates, extending the general applicability of ^1^H *R*_1ρ_ to probe dynamics at natural abundance. Artifacts were found to be negligible for neighbouring dipolar-coupled protons, >3 Å apart, and a concept for identification for protons less than 3Å is provided. This approach revealed a previously hidden, second excited state (ES2) in DNA base-pairing that extends the well-established Watson-Crick-Franklin (WCF) ground state (GS) – Hoogsteen (HG) equilibrium. A structural model for ES2 is proposed based on evidence from ^1^H *R*_1ρ_ RD, trapping via DNA modifications, metadynamics simulations, and DFT-based chemical shift calculations. ES2 was stabilised by the anticancer drug Actinomycin D, providing direct experimental evidence that small molecule can remodel conformational landscape of DNA. Together, these results demonstrate both a methodological advance by establishing reliable conditions for ^1^H *R*_1ρ_ studies, and a mechanistic discovery of a drug-stabilized intermediate in DNA base-pairing dynamics.

## Introduction

Conformational dynamics in biomolecules play a crucial role in defining their biological function^1–3^. In these biomolecules, microsecond to millisecond time scale dynamics are present between an energetically favourable ground state (GS) and a higher energy, excited state (ES)^4–6^. ES conformations have been reported to have important roles in nucleic acids, e.g. microRNA processing^7,8^, targeting^1^, DNA base repair^9^, and HIV activation^10^. Due to their low population (typically < 2%)^4,5^, ES are challenging to characterise using either X-ray crystallography or cryo-electron microscopy. NMR spectroscopy, however, has proven to provide atomic-resolution structural and dynamical information about these ES^6,10–12^.

Measuring the longitudinal relaxation rate in the rotating frame (*R*_1ρ_) and its dispersion with respect to the applied spin-lock field strength (ω_SL_; on-resonance) and spin-lock offset relative to the resonance of interest (Ω_SL_; off-resonance) has been the method of choice to identify such ES^6^. *R*_1ρ_ relaxation dispersion (RD) quantifies the conformational exchange contribution (*R*_ex_) to the transverse relaxation rate (*R*_2_) of the resonance under study^6,13^. It can probe exchange rates between 50 Hz to 50 kHz and provide access to the chemical shift of the ES^6,14–16^. This chemical shift information allows modelling the structure of an ES^12^.

The ES conformation typically observed in DNA is the Hoogsteen (HG) base-pair, which is crucial in processes like DNA damage repair^17,18^, and recognition of DNA by transcription factors^19,20^. In an HG base-pair, the purine nucleobase adopts a *syn* conformation around the glycosidic bond (C1’–N9) rather than *anti* as in a Watson-Crick-Franklin (WCF) (Figure 1a and1b)^21,22^. ^13^C and ^15^N RD experiments have been employed to characterize the HG ES in DNA^2,3,6,9,11,23–27^. Recently, a ^1^H high-power chemical exchange saturation transfer (CEST) experiment was developed to study WCF – HG dynamics^21^ in a model DNA oligonucleotide, A_2_ DNA (Figure 1a)^3,21^. The exchange rates that can be probed with these methods, however, are limited to *k*_ex_ up to 4–5 kHz^6^. The ^1^H *R*_1ρ_ experiment expands the accessible exchange rate to tens of kHz, is more sensitive compared to CEST^14,15^ and can be readily applied to systems at natural isotopic abundance. Additionally, ^1^H chemical shifts are particularly sensitive towards identifying non-canonical base-pairs conformation in nucleic acids^28^.

**Figure 1.**
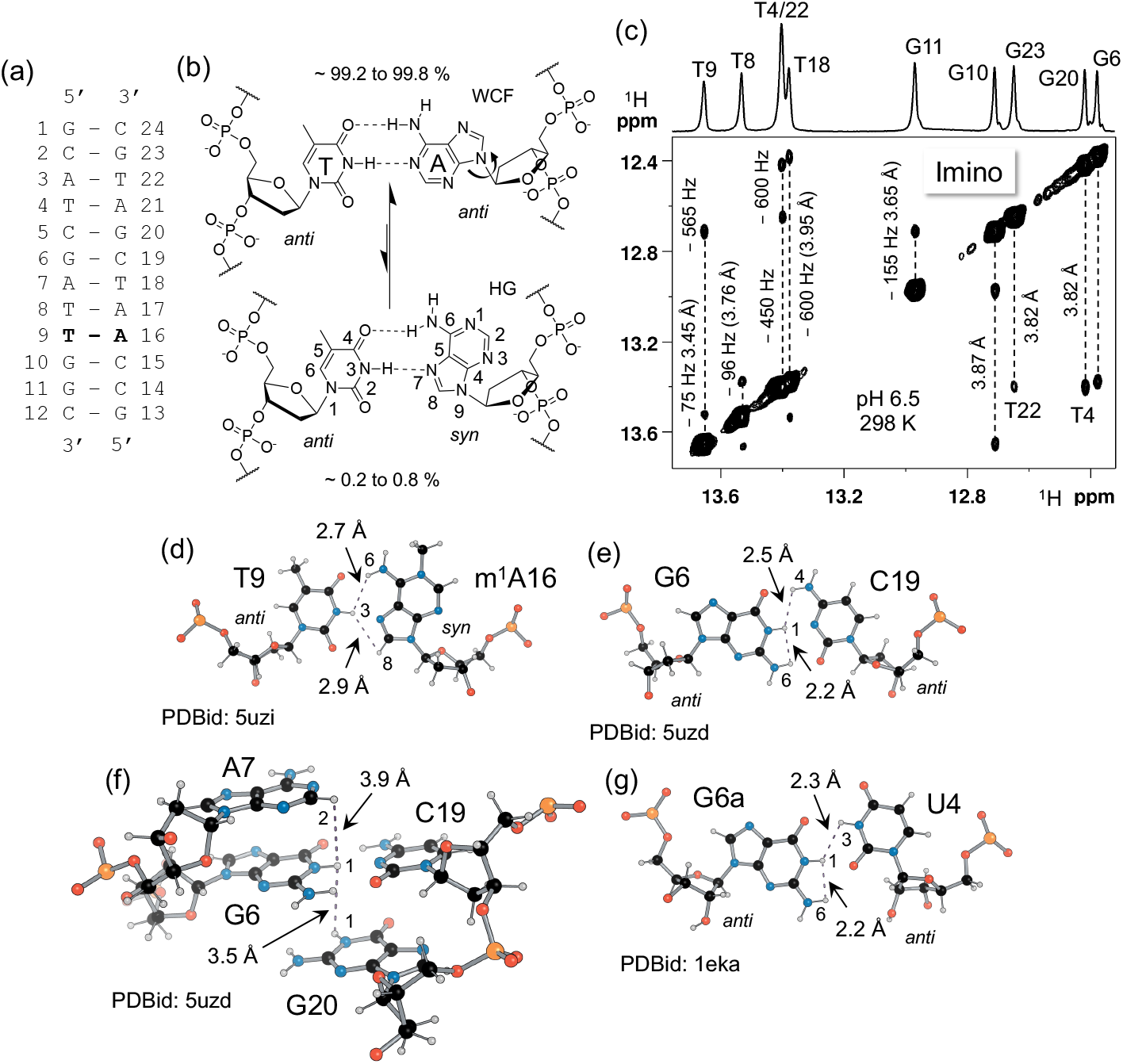
Model system used in this study: (a) A_2_ DNA sequence^3^ with the base-pair T9–A16 marked in bold, where a representative WCF–HG dynamics was measured. (b) WCF–HG conformations for the T9–A16 base-pair with the reported populations and *anti-*to-*syn* transition for A16^3,21,24^. (c) Selective imino NOESY^30^ spectrum of A_2_ DNA. The 1D projection is shown at the top of the spectrum with the resonance assignment^21,24^. The position of the cross-peaks (in Hz) and the corresponding average distance from the 10 minimum energy structures (in Å) in PDB 5uzd^29^ are denoted. (d-g) Representative NMR-derived intra- and inter-base-pair proton–proton distances that may contribute to ^1^H *R*_1ρ_ relaxation rates: (d) T9-m^1^A16 HG base-pair in A_6_ DNA, PDBid 5uzi^29^, (e) G6:C19 WCF base-pair in A_2_ DNA, PDBid 5uzd^29^, (f) Base-pair steps from G6-C19 to the neighbouring A7 and G20 in A_2_ DNA, PDBid 5uzd^29^ and (g) U4-G6 wobble base-pair in r(GAGUGCUC)_2_ RNA, PDBid 1eka^31^. Atomic position numbers are provided for protons involved in hydrogen bonds.

^1^H *R*_1ρ_ RD for a given proton might, however, have contributions from neighbouring protons due to cross-relaxation that could be mistaken for additional conformational exchange^14^. An NMR structure ensemble of A_2_ DNA^29^ indicates ^N^H–^N^H distances in the range of 3.4–3.9 Å between consecutive base-pairs (Figure 1c). Distances from ^N^H to other protons within a base-pair in typical nucleic acid geometries range from 2.2 Å to 2.9 Å in various structural contexts in both DNA and RNA (Figure 1d–g). These distances might pose a challenge to obtain reliable exchange parameters from ^1^H *R*_1ρ_ experiments on A_2_ DNA.

This study shows that cross-relaxation between ^N^H protons at typical inter-base-pair distances as described above has negligible impact on *R*_1ρ_ RD profiles, which consequently can be analysed using well-established models for conformational exchange^6,32–35^. A significant effect is seen at shorter interproton distances (Figure 1d, 1e, and 1g), which can be mitigated by prudent choice of spin-lock strengths and offsets in the *R*_1ρ_ experiment. A thorough theoretical description of cross-relaxation effects on ^1^H *R*_1ρ_ relaxation dispersion using Bloch-McConnell (BM) matrix propagation^36^ is discussed under different exchange scenarios. In A_2_ DNA, this allowed the detection of a previously unreported excited state, ES2, involved in the WCF–HG equilibrium. To explore the structural implications of ES2, the HG-forming adenine in the central T9-A16 base-pair was chemically modified to obtain chemical shift fingerprints of different base-pair conformations^12^. This ES2 is also shown to be promoted in presence of an anti-tumour drug Actinomycin D. The well-tempered parallel bias metadynamics simulation^37–39^ in combination with chemical shift calculation a structural model for the ES2 is proposed. Overall, the discovery of ES2 offers a deeper understanding of the energy landscape of the WCF–HG transition, which may ultimately provide significant insights into its role in DNA biochemistry.

## Results and Discussions

### Effects of cross-relaxation on ^1^H R_1ρ_

To analyse the effect of cross-relaxation on *R*_1ρ_ RD profiles in the R1ρ experiment (Figure 2a), a model involving a two-state exchange between a GS and ES in the presence of neighbouring dipolar coupled proton (^1^H_dip_) was considered (Figure 2b). The ^1^H_dip_ was positioned at distances of *r*_i_ and *r*_j_ relative to the GS and ES, respectively, which can be varied to consider multiple scenarios. Transverse (*R*_2_) and longitudinal (*R*_1_) auto-relaxation rates, together with chemical exchange, are described by the Bloch-McConnel (BM) equations (equations S1–S16)^6,32–34,40^. These can be expanded to include transverse (μ) and longitudinal (σ) cross-relaxation rates induced by ^1^H_dip_ (equation S17–S18)^36^. The resulting BM matrix can be propagated over the pulse sequence “ramp (φ = −y) – spin-lock strength (ω_SL_) (φ = x) – ramp (φ = y)” (Figure 2a), the main component of the *R*_1ρ_ RD experiment^14,15^. The initial magnetization vector aligned along the z-axis was weighted using the populations of the GS (*p*_GS_) and ES (*p*_ES_) as specified during the simulation. For simplicity, the ^1^H_dip_ magnetization was not weighted since it is assumed to be a static proton not undergoing any chemical exchange. Isotropic spectral density function, J(ω) (equation S7), was calculated with a rotational correlation time (τ_c_) of 5.1 ns based on the size of the A_2_ DNA duplex^29^.

**Figure 2.**
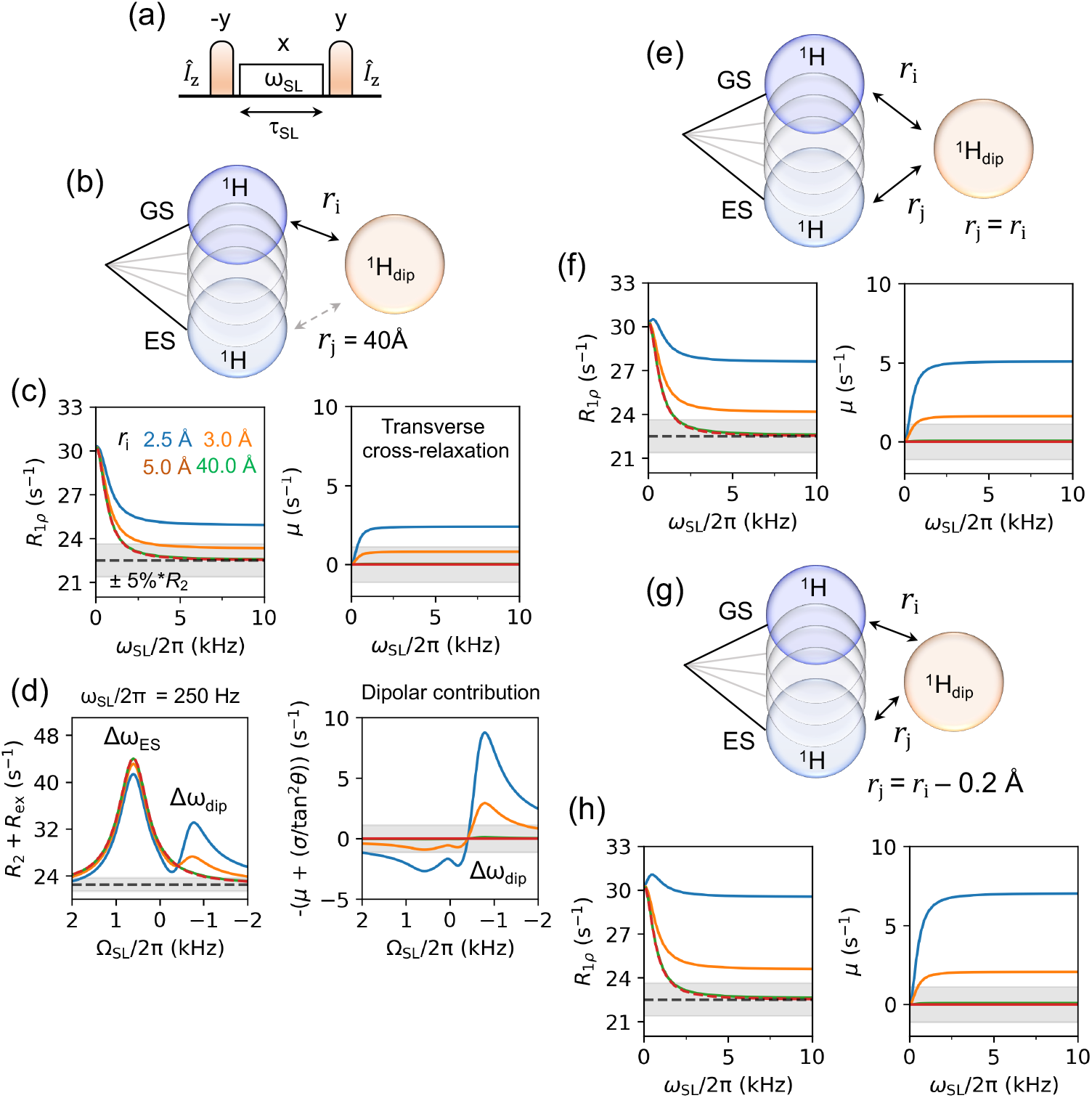
Effects of cross-relaxation on ^1^H *R*_1ρ_ experiments. **(a)** *R*_1ρ_ pulse sequence used in simulations: z magnetisation on spin *I* (*I*_z_) is rotated to the zx plane by a *−y* pulse, spin-locked with strength ω_SL_ for τ_SL_, and returned to z-axis with a *y* pulse. **(b)** Two-site chemical-exchange model between ground state (GS) and excited state (ES) (blue circles), with a dipolar-coupled proton, ^1^H_dip_ (light brown). Distances between ^1^H_dip_ and GS is denoted as *r*_i_ and *r*_j_, respectively where *r*_j_ = 40Å representing scenario 1. (c) On-resonance profile (left) for *r*_i_ = 2.5, 3.0, 5.0 and 40Å (blue, orange, brown, and green) depicting that at for *r*_i_ ≥ 3 Å, cross-relaxation effects remain within ± 5% (grey region, representing typical experimental error) of the *R*_2_ rate, ensuring accurate extraction of exchange parameters. Transverse cross-relaxation (μ) rate contribution (right) to the on-resonance *R*_1ρ_ profile shows the same trend. (d) Off-resonance *R*_2_ + *R*_ex_ profile (left) for scenario 1 at ω_SL_/2π = 250 Hz with Δω_ES_ and Δω_dip_ depicted in the plot. This shows that the dipolar proton has a response at its Δω_dip_ at ≤ 3Å. The dipolar contribution on the off-resonance profile (right) shows that there is a substantial contribution on the response from ES (Δω_ES_) at distances < 3Å thereby complicating the extraction of reliable exchange parameters. (e, g) Two-state exchange model where *r*_j_ = *r*_i_ or *r*_j_ = *r*_i_ – 0.2 Å representing scenario 2 and 3 respectively. (f, h) On-resonance profile (left) and contribution from μ (right) shows that cross-relaxation in the ES amplifies *R*_2_ contribution (*μ*) more in scenario 2 and 3 than in scenario 1 where the distance to the ^1^H_dip_ must be > 3Å to be within ± 5% error range of *R*_2_. Parameters used for the simulations are *k*_ex_ = 2 kHz, *p*_ES_ = 0.5%, τ_c_ = 5.1 ns, *R*_1GS_ = *R*_1ES_ = *R*_1dip_ = 2.5 s^-1^, *R*_2GS_ = *R*_2ES_ = *R*_2dip_ = 22.5 s^-1^, Δω_ES_/2π = +600 Hz and Δω_dip_/2π = -600 Hz.

*R*_1ρ_ RD experiments can be performed by varying two parameters: (a) the spin-lock strength (ω_SL_), which is positioned on-resonance at the observed chemical shift, and (b) the position of ω_SL_ known as the offset (Ω_SL_) which is varied by approximately ±4 × ω_SL_ relative to the observed chemical shift, to sample the resonance position of the ES^6^. The Z-magnetization (M_z_) under increasing duration of ω_SL_ (τ_SL_) decays monoexponentially with the rate constant 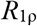.

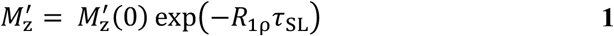

In the absence of 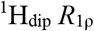 is represented as^32,33^

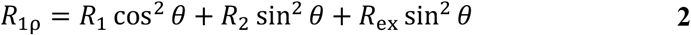

Here, *R*_1_ and *R*_2_ are the intrinsic longitudinal and transversal auto-relaxation rates, *θ* is the angle of the effective field from the Z-axis (*θ* = arctan (ω_SL_/Ω_SL_)) and *R*_ex_ is the chemical exchange contribution to the *R*_2_. Equation 2 can be extended to account for the effects of μ and σ as (equation S17–S19)

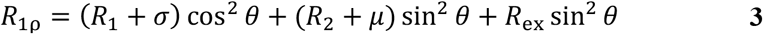

On-resonance, where θ = 90°, the 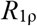 rate from equation 3 approaches *R*_2_ + μ in the high spin-lock limit while off-resonance, when Ω_SL_ >> ω_SL,_ it approaches *R*_1_ + σ. Within this framework, the following scenarios were considered to discuss the effect of cross-relaxation on both on- and off-resonance experiments using common exchange parameters of *k*_ex_ = 2 kHz, *p*_ES_ = 0.5%, τ_c_ = 5.1 ns, *R*_1GS_ = *R*_1ES_ = *R*_1dip_ = 2.5 s^-1^, *R*_2GS_ = *R*_2ES_ = *R*_2dip_ = 22.5 s^-1^, Δω_ES_/2π = +600 Hz and Δω_dip_/2π = -600 Hz:

#### Scenario 1: Cross-relaxation only in the GS and not the ES

The distance *r*_i_ (^1^H_dip_ to GS proton) was varied from 2 Å to 4 Å, while cross-relaxation in the ES was neglected by setting *r*_j_ (^1^H_dip_ to ES proton) to a sufficiently large value (40 Å). This approach models a dynamic nucleic acid system, such as a base flipping out of the helical axis in the ES, which increases the distance between the proton of interest and ^1^H_dip_ compared to the GS. Simulated on-resonance *R*_1ρ_ (Figure 2c) reveals an increased μ contribution when *r*_i_ < 3 Å, corresponding to the ^N^H1–^N^H3 distance (2.3 Å) in GU/GT wobble base-pairs (Figure 1g). At distances *r*_i_ < 2.5 Å–typical of geminal protons such as H5’, H5’’ and H2’, H2’’ in the deoxyribose sugar and aromatic H6, H5 protons–*R*_1ρ_ initially rises at ω_SL_ < 1 kHz and approaches the *R*_2_ + μ in the high spin-lock limit (Figure S1a). At *r*_i_ ≥ 3 Å, which represents inter-imino proton distances in canonical B-form DNA and A-form RNA, the μ contribution remains minimal, contributing ≤ 5% to *R*_2_ (Figure 2c).

The simulated off-resonance profile (Figure 2d) demonstrates that for Ω_SL_ >> ω_SL_, *R*_1ρ_ approaches *R*_1_ + σ. At *r*_i_ ≤ 2.2 Å, this sum becomes negative due to the dominant contribution of σ, leading to an exponential increase in Z-magnetization (Figure S1b). This magnetization buildup complicates the estimation of *R*_2_ + *R*_ex_ and necessitates an approximate analytical solution beyond the scope of this study. For *r*_i_ between 2.2 and 2.5 Å and assuming Δω_dip_ = -600 Hz (1 ppm at a 600 MHz ^1^H Larmour frequency used in this work), the *R*_2_ + *R*_ex_ profile exhibits a response from ^1^H_dip_, characterized by a shift in local maximum of *R*_2_ + *R*_ex_ as ω_SL_ increases (Figure 2d, Figure S1c). This behaviour, which distinguishes it from ES to GS exchange, diminishes at *r*_i_ ≥ 3 Å. In contrast, the intra-base-pair protons, such as amino (-NH_2_), aromatic (^C^H2 and ^C^H8), and imino (^N^H3/H1) protons at distances *r*_i_ ≤ 3Å (Figure 1d, 1e and 1g) do not exhibit this effect due to their large chemical shift differences of 2.1 to 2.7 kHz (-3.5 to -4.5 ppm), which lie beyond the conventionally probed Ω_SL_ range (±4x ω_SL_). However, if Ω_SL_ is near Δω_dip_, a response is observed (Figure S1d).

Simulations varying *k*_ex_ and Δω_dip_ (Figure S1e – S1h) indicate minimal cross-relaxation effects at *r*_i_ ≥ 3 Å and |Ω_SL_| < |Δω_dip_|. The impact of Δω_dip_ on the ^1^H *R*_1ρ_ experiment was also evaluated. For *r*_i_ = 2.5 Å, an increase in on-resonance *R*_1ρ_ rate is observed when Δω_dip_ approaches the GS resonance (Figure S2a-c). When Δω_dip_ and Δω_ES_ are in the same direction, accurate extraction of chemical exchange parameters becomes challenging due to difficulty in disentangling the contribution of exchange and cross-relaxation to *R*_1ρ_ (Figure S2b). For *r*_i_ ≥ 3.0 Å, these effects are mitigated, enabling reliable extraction of exchange parameters.

#### Scenario 2 and 3: Cross-relaxation equal in both GS and ES or is larger in ES

In this scenario, chemical exchange involves an ES conformer susceptible to cross-relaxation, where the neighbouring ^1^H_dip_ is positioned such that: (a) the ^1^H_dip_ is equidistant to the proton of interest in both GS and ES (Figure 2e), or (b) the ^1^H_dip_ is closer to the proton of interest in the ES compared to the GS (example: *r*_j_ = *r*_i_ – Å) (Figure 2g). Simulations of on-resonance profiles reveal that the cross-relaxation contribution increases as the distance *r*_j_ decreases relative to *r*_i_. The *R*_1ρ_ rates approach *R*_2_ + μ at *r*_i_ = 3.3 Å (within ± 5% error), compared to 3 Å in scenario 1 (Figure 2f, 2h). For *r*_i_ < 2.4 Å, *R*_2_ + μ exceeds *R*_ex_. This enhanced cross-relaxation also amplifies the exponential rise observed in the off-resonance profile under this scenario (Figure S2d-g). Consistent with scenario 1, cross-relaxation effects are minimal at distances *r*_i_ ≥ 3Å.

As a result, ^1^H *R*_1ρ_ provides a valuable tool for studying conformational exchange involving nucleobase protons, provided Δω_ES_ lies within and Δω_dip_ remains outside the probed Ω_SL_ range. Prior knowledge of neighbouring protons and chemical shifts can help in designing specifically labelled sample and estimate cross-relaxation effects.

### ^1^HR_1ρ_ RD reveals third state in WCF–HG transition

^13^C and ^15^N *R*_1ρ_ as well as high-power ^1^H CEST experiments on A_2_ DNA have demonstrated that WCF–HG dynamics follow a two-state exchange model with an exchange rate of 3 to 4 kHz and HG population of 0.2 to 0.8%^3,21,24^. Motivated by the above results of manageable cross-relaxation effects, ^1^H *R*_1ρ_ experiments were conducted to study the WCF–HG transition in A_2_ DNA. The NOESY spectrum of the imino ^N^H region shows that the dipolar-coupled T9 ^N^H3 – T8 ^N^H3 and T9 ^N^H3 – G10 ^N^H1 pairs are at chemical shifts of −75 Hz (-0.12 ppm) and −565 Hz (-0.94 ppm), respectively, relative to T9 ^N^H3 (Figure 1c). Given their ∼ 3.9 Å distance^29^, these interactions minimally influence the on- and off-resonance ^1^H *R*_1ρ_ rates (Figure 2c and 2d).

^1^H *R*_1ρ_ data for T9 ^N^H3 revealed two conformational exchange contributions (Figure 3a, S3a, Table S1) modelled using a triangular three-state exchange topology (equation S20) and statistically preferred over all other tested models by F-test, Akaike Information Criterion (AICc), and Bayesian Information Criterion (BIC)^6,41^ (Supporting Excel file). Comparing with the previous reported chemical shifts^21^, the conformation at Δω = −593 ± 10 Hz (-0.99 ± 0.02 ppm) corresponds to the HG state (Δω_HG_), with exchange rate (*k*_ex, WCF ⇋ HG_) = 2.7 ± 0.15 kHz and population (*p*_HG_) = 0.6 ± 0.01% (Figure 3a and 3b, Table S1). Additionally, a second excited state (ES2) at Δω_ES2_ = +288 ± 9 Hz (+0.48 ± 0.01 ppm) with *p*_ES2_ = 0.9 ± 0.1% was observed. The exchange rates with ES2 are *k*_ex, WCF ⇋ ES2_ = 0.5 ± 0.07 kHz and *k*_ex, HG ⇋ ES2_ = 3.2 ± 0.15 kHz (Figure 3b, Table S1). This ES2 represents an intermediate during the WCF-HG transition in A_2_ DNA. Furthermore, the system displays a temperature-dependent change in topology. A linear topology dominated at 283, 288, and 308 K, while a triangular topology was observed at 293, 298, and 303 K (Figure S4). This behaviour, coupled with the non-linear van’t Hoff plot previously observed^3^, likely arises from the presence of ES2, thereby making the derivation of reliable thermodynamic parameters challenging.

**Figure 3.**
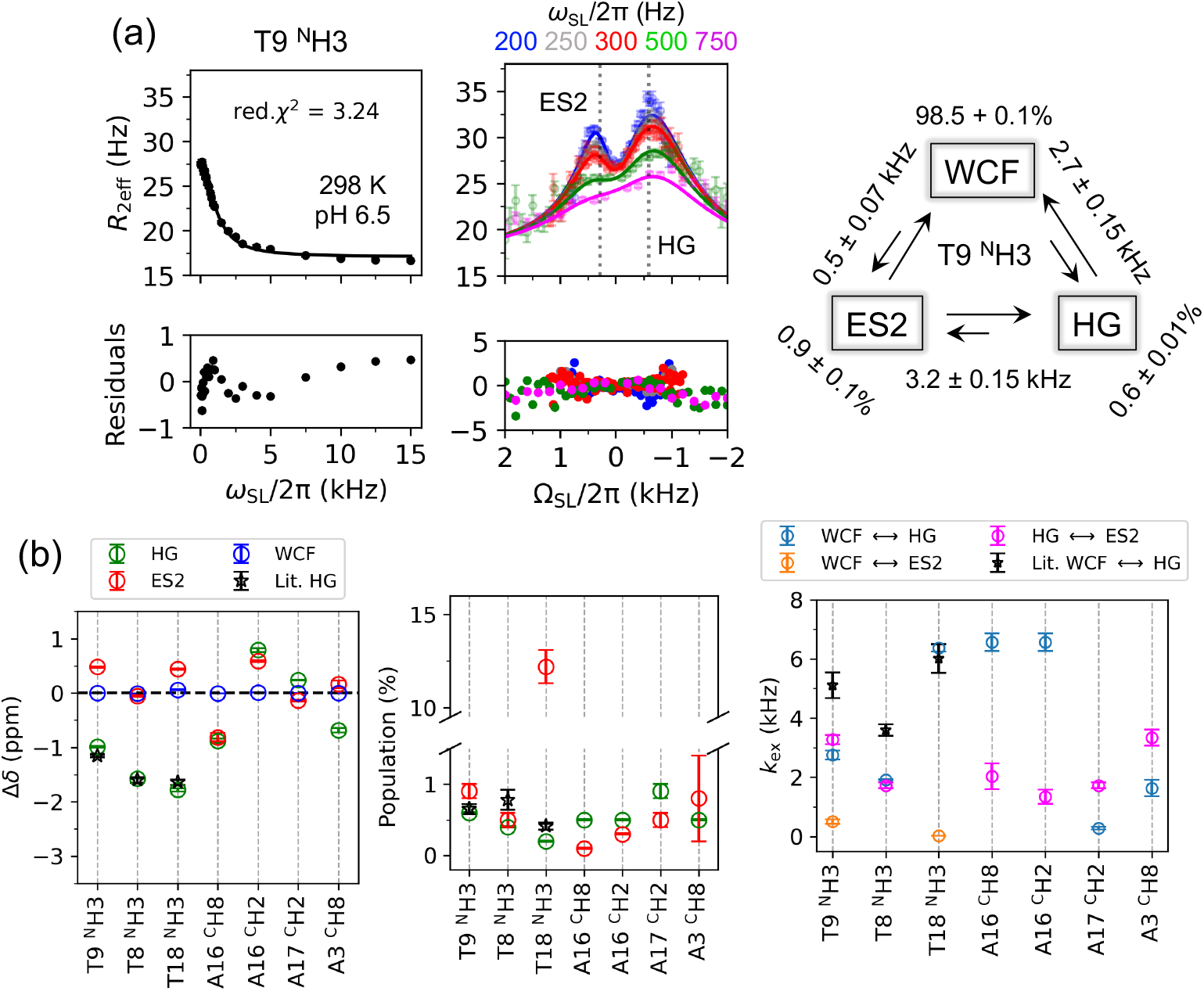
^1^H *R*_1ρ_ RD reveals ES2 in WCF – HG dynamics in A_2_ DNA. (a) On-resonance (left) and off-resonance (right) *R*_2eff_ plots for T9 ^N^H3 at 298 K and pH 6.5 with a three-state exchange fit. Solid lines represent the fits associated with the fit parameters (Table S1), and dashed lines show relative chemical shifts of HG and ES2. Exchange rates and populations for each conformer are shown in the schematic representation of the best fitted three-state triangular exchange model. (b) Chemical shifts for WCF (blue), HG (green) and ES2 (red) are shown for each studied atom. Δω_HG_ and *p*_HG_ from ^1^H *R*_1ρ_ of ^N^H3 in T9, T8, and T18 are consistent with the literature data from high-power ^1^H CEST^21^ (black star). This confirms ES2 as an intermediate state during the WCF – HG transition.

^1^H *R*_1ρ_ experiments on the base-pairing partner of T9, A16, which undergoes the *anti*–to-*syn* transition to form the HG conformer, further confirmed the presence of ES2. Non-exchangeable aromatic protons ^C^H2 and ^C^H8 were globally fitted to a three-state linear topology (Figure S3b, S3c and S3e, Table S1, Supporting Excel file), sharing *k*_ex, WCF ⇋ HG_, and *p*_HG_ values. However, their sensitivity to WCF–ES2 transitions was limited, resulting in higher uncertainty in the estimation of the ES2 population. Consequently, a global fit combining T9 and A16 ^1^H *R*_1ρ_ data was not feasible due to differing exchange rates and populations. Other protons in the neighbouring T:A base-pairs (T8 ^N^H3, T18 ^N^H3, and A17 ^C^H2) also indicated the presence of an ES2 (Figure S3). T8 ^N^H3 and A17 ^C^H2, ES2 were possible to be globally fitted to a linear three-state exchange model with a shared *k*_ex, HG ⇋ ES2_, and *p*_ES2_, without resolved WCF–ES2 contributions (Figure 3b, Table S1). T18 ^N^H3 showed three-state star-like exchange where only the WCF–ES2 transition was resolved, with no HG–ES2 transition detected. The reason for the associated comparatively high *p*_ES2_ (12.2 ± 0.9%) and low *k*_ex, WCF ⇋ ES2_, (24 ± 0.7 s^-1^) could not be explained with the current dataset. This suggests that certain triangular exchange components may remain unresolved in some situations. Nevertheless, the observed *p*_HG_ and Δω_HG_ exchange parameters for T9, T8, and T18 ^N^H3 match the previous report using high-power ^1^H CEST^21^ (Figure 3b), while the difference in *k*_ex, WCF ⇋ HG_ can be attributed to different pH values of the sample used in the present study. Overall, these findings establish that ES2 is a consistent intermediate state within T:A base-pairs in A_2_ DNA, reflecting the complex dynamics of WCF-HG transitions.

### Structural information of ES2 by chemical modification of A16

To gain insight into the conformation of ES2, three A_2_ DNA constructs with chemically modified nucleobases at A16 were prepared (Figure 4a, Tables S2, S3): (i) Purine (A_2_ P16) substitution to eliminate the amino group that hydrogen-bonds with O4 of T9^42^, reducing base-pair strength in both WCF and HG conformations; (ii) 7-deazaadenine substitution (A_2_ c^7^A16), which removes the N7 hydrogen bond acceptor, favouring the WCF conformation by sterically interfering with HG base-pair formation^24^; and (iii) 1-methyladenine substitution (A_2_ m^1^A16), where methylation of N1 promotes formation of the HG conformation by blocking the WCF edge of A^3^. Chemical shift perturbations (CSPs) of imino protons due to these modifications are localized primarily to the T9-A16 base-pair and its adjacent ±1 neighbours (Figure 4b). NOESY and SOFAST HMQC spectra confirmed T9 WCF base-pairing with A_2_ P16 and A_2_ c^7^A16, and HG base-pairing with A_2_ m^1^A16 (Figure S5-S7), consistent with prior reports^24,25,29,42,43^. Imino ^N^H3/1 chemical shifts showed that in the GS conformation, T9 ^N^H3 of both A_2_ P16 and A_2_ m^1^A16 has a lower chemical shift, while in A_2_ c^7^A16 it has a higher chemical shift relative to the wild-type DNA (A_2_ wt) (Figure 4c). This deshielding in A_2_ c^7^A16 was attributed to the altered electrostatic properties of the c^7^A modification, increased solvent accessibility, and changes in stacking interactions^24,43^. Coincidently, the T9 ^N^H3 shift in A_2_ c^7^A16 matched that of ES2 in A_2_-wt (Figure 4c), a state with a lower base-pairing stability.

**Figure 4.**
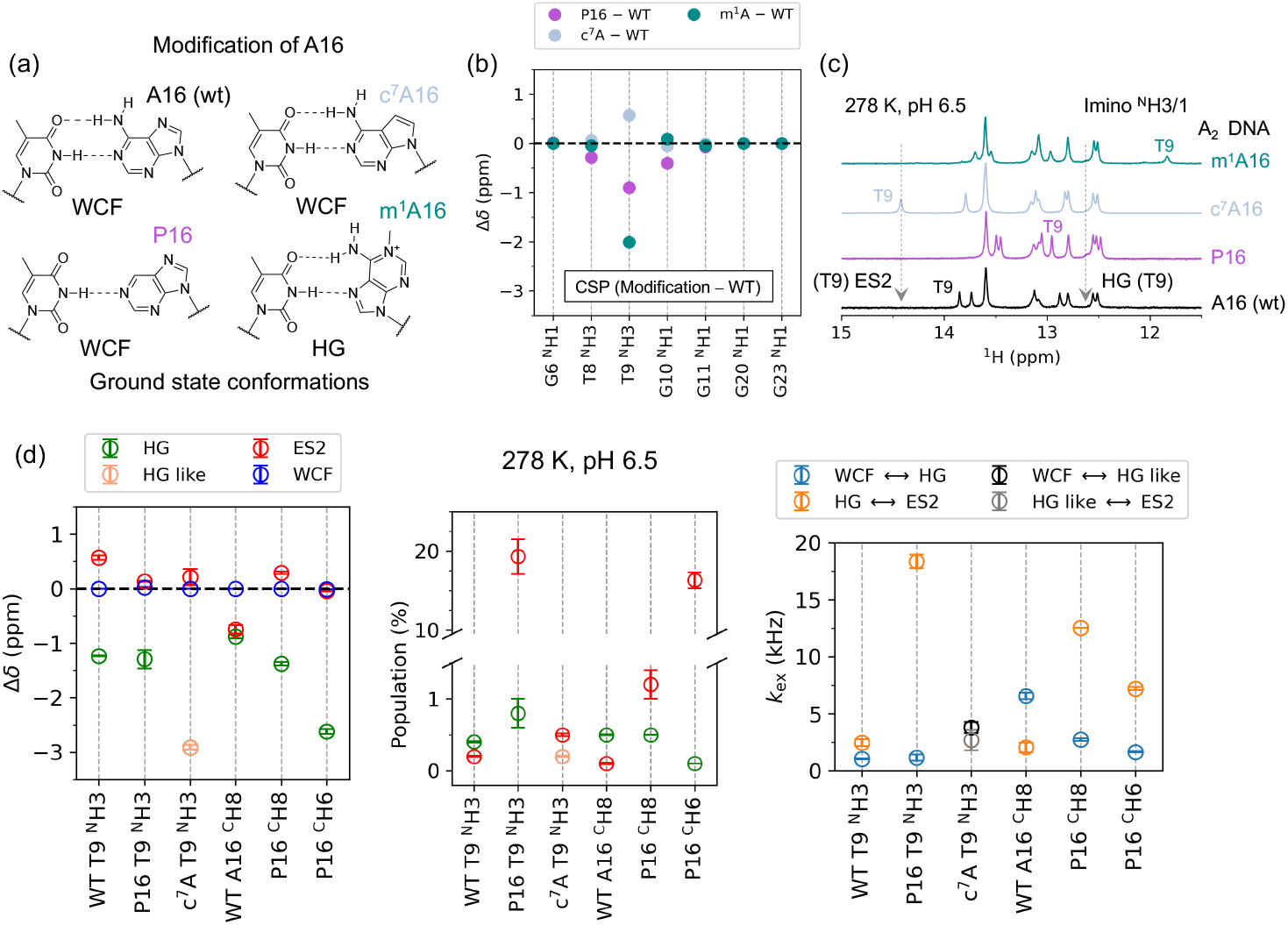
Effects of A16 modification on WCF – HG – ES2 dynamics. (a) Chemical structures of unmodified (A_2_-wt) and modified A16 nucleobases (A_2_-P16, -c^7^A16, and - m^1^A16) used to modulate the WCF-HG exchange equilibrium in A_2_ DNA. (b) Chemical shift perturbations (CSPs) of imino protons in modified A_2_ DNA constructs show localized effects near the site of modification (T9–A16 base-pair), with minimal influence on adjacent base-pairs and no detectable perturbation beyond ±1 neighbours. (c) ^1^H imino spectra at 278 K and pH 6.5 for A_2_-wt (black), A_2_-P16 (magenta), A_2_-c^7^A16 (steel blue), and A_2_-m^1^A16 (dark green), highlighting the position of the T9 ^N^H3 resonance. Arrows indicate the chemical shift corresponding to the minor ES2 and HG conformations as identified in T9 of A_2_-wt by ^1^H *R*_1ρ_ experiments. (d) Kinetic parameters derived from ^1^H *R*_1ρ_ experiments for T9 ^N^H3 in A_2_-P16 and A_2_-c^7^A16, as well as aromatic protons (^C^H6 and ^C^H6) of the modified P16 base. Comparison of Δδ (ppm) with A_2_-wt confirms the presence of both HG (green) and ES2 (red) in A_2_-P16. Notably, the population of ES2 and the associated exchange rate *k*_ex(HG – ES2)_, (orange) are elevated in A_2_-P16, while *k*_ex(WCF – HG)_, (light blue) remains unchanged. In A_2_-c^7^A16, a HG-like state (light orange) with respective exchange rates (grey and black) is observed despite prior assumption^24^ that steric hindrance between the A16 ^C^H7 and T9 ^N^H3 would prevent such a conformation.

Using ^1^H *R*_1ρ_ RD at 278 K in A_2_ P16, the T9 ^N^H3, P16 ^C^H2 and P16 ^C^H8 signals fitted a three-state linear exchange model (Figure 4d, S8, supporting Excel file). The fitted Δω_HG_ conformation has the same sign as the HG-trapped A_2_ m^1^A16 DNA, while the Δω_ES2_ followed the sign observed in A_2_ wt (Figure 3b), supporting this assignment. A_2_ P16 showed increased *k*_ex, HG ⇋ ES2_ (18 ± 0.6 kHz) and *p*_ES2_ (19.3 ± 2.2 %) (Figure 4d, Table S4 and S5) compared to A_2_ wt (Figure S4a and supporting Excel file). This suggests that the increased flexibility of the T9-P16 base-pair enhances ES2 sampling. Similar exchange patterns observed for P16 ^C^H8 and P16 ^C^H2 (Figure 4d, S8, supporting Excel file) indicate that the findings are not confounded by water exchange on the imino proton, despite lower relative stability of the T9-P16 base-pair^44^. The 20-fold increase in *p*_ES2_ and 10-fold increase in *k*_ex, HG ⇋ ES2_ for A_2_ P16 with respect to A_2_ wt underscore the role of the amino group in limiting ES2 sampling.

In A_2_-c^7^A16, ^1^H *R*_1ρ_ RD at 278 K for T9 ^N^H3 also revealed a three-state linear exchange (Figure 4d, S8) with one excited state at Δω = −1748 ± 27 Hz (-2.91 ± 0.04 ppm), consistent with the HG conformation observed in other DNA duplexes^3,21,45^. This is intriguing since the c^7^A modification sterically disfavours the HG conformation, suggesting this is an HG-like state that requires further investigation. The other excited state exhibited Δω = 127 ± 89 Hz (+0.21 ± 0.13 ppm), resembling the ES2 in A_2_ wt. The exchange parameters for this ES2-like state in A_2_-c^7^A16 were comparable to those in A_2_-wt (Figure S4a, supporting Excel file), indicating that N7 does not influence the ES2 conformation as significantly as the N6 amino group, as shown in A_2_-P16. Notably, this exchange in c^7^A-modified DNA was undetected in earlier ^13^C and ^15^N *R*_1ρ_ studies^24^, highlighting the enhanced sensitivity of ^1^H *R*_1ρ_ RD.

For A_2_-m^1^A16 DNA, ^1^H *R*_1ρ_ RD on T9 ^N^H3 at 278 K exhibited a dispersive on-resonance profile, indicative of a minor conformation with *k*_ex_ > 25 kHz. The rapid exchange precluded precise estimation of population and Δω values from off-resonance experiments (Figure S7e, Table S4).

### Actinomycin D binding promotes ES2 conformation

To investigate the relevance of ES2 in presence of a known DNA-binding drug, A_2_ wt DNA was treated with the cytostatic compound Actinomycin D (ActD). Although the A_2_ DNA does not contain the canonical GpC binding site for ActD^46^, CSP analysis suggests the presence of two binding modes (A and B) at 1:1 concentration (1.0 mM each ActD and A_2_ DNA) and 298 K (Figure 5a and 5b). These binding sites are localized around T9, G10 and G11, resembling various non-canonical binding sites such as the GpG and T(G)_n_T sites reported in the literature^47–53^. Overlaying the chemical shift data of ES2 in A_2_ wt onto these CSPs indicates that in binding mode B, the observed chemical shifts of ES2 for T9 ^N^H3 and A16 ^C^H2 align most closely (Figure 5b and S9), suggesting that this state is stabilized or promoted, indicating a potential role of ES2 in drug interactions.

**Figure 5.**
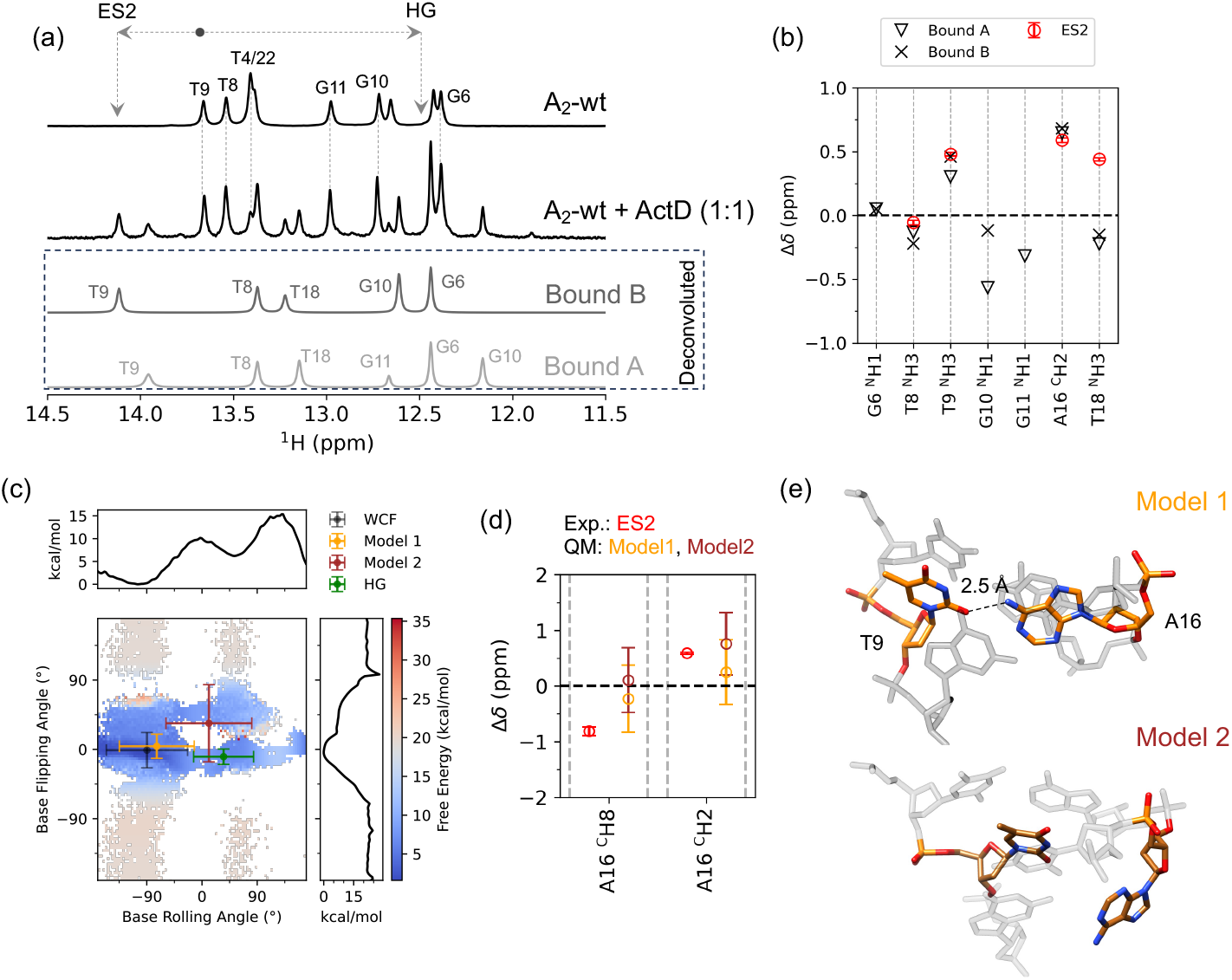
Drug interaction and structural models of ES2. (a) ^1^H imino spectra of free and Actinomycin D-bound A_2_ DNA (1:1 molar ratio) at 298 K and pH 6.5 reveal two distinct bound states: Bound A (light grey) and Bound B (dark grey). Deconvoluted spectra for each bound state are shown below. Arrows indicate the chemical shifts corresponding to the ES2 and HG conformations for T9 ^N^H3 (black dot) in A_2_ wt. Notably, the T9 ^N^H3 resonance in Bound B closely matches the ES2 position. (b) Chemical shift difference (Δδ, ppm) between the ActD-bound (triangles: Bound A; crosses: Bound B) and free A_2_ DNA are overlaid with the experimental Δδ associated with ES2 (red). The alignment of chemical shifts for T9 ^N^H3, T8 ^N^H3, and A16 ^C^H2 between Bound B and ES2 supports the hypothesis that ES2 may contribute to drug binding. (c) Reweighted 2D free energy surface (FES) from metadynamics simulations plotted as a function of base-flipping and base-rolling angle (χ-dihedral). Cluster centroids derived from chemical shift-based agglomerative clustering are overlaid: WCF (black) and HG (green) correspond to the expected minima. Model 1 (orange) and Model 2 (brown) map clusters in distinct regions, where Model 1 aligns with a local low-energy region, while Model 2 occupies a higher-energy region. This trend is consistent across multiple collective variables CV projections (Figure S12), suggesting Model 1 as a more plausible structural candidate for ES2. One-dimensional projections of the FES are shown along the top and right axes. (d) Relative chemical shift (Δδ, ppm) for non-exchangeable A16 ^C^H8 and A16 ^C^H2 aromatic protons from experimental ES2 (red) and calculated Model 1 (orange), and Model 2 (brown). Error bars represent one standard deviation across cluster frames. Both models exhibit chemical shift values consistent with the experimental ES2. (e) Representative minimum-energy structures from the Model 1 and Model 2 clusters. Model 1 features a hydrogen bond between the A16 amino group and T9 O2 (dashed line with heavy atom distance of 2.5Å), while Model 2 exhibits partial flipping of A16 out of the helical axis.

### Structural models for ES2

Molecular dynamics (MD) simulations were performed to investigate the WCF and HG conformations of the T9-A16 base-pair in A_2_-wt, A_2_-P16 and A_2_-c^7^A16 DNA constructs. The WCF conformation was modelled based on a reported NMR structure (PDB ID: 5UZD)^29^, while the HG conformation was generated by rotating the χ-dihedral angle of A16 by 180° followed by energy minimization. Parameters for the modified nucleotides P16 and c^7^A were derived using standard two-stage restrained electrostatic potential (RESP) fitting of *N*^9^-methylated nucleobases with geometries optimized at the HF/6-31G^*^ level of theory^54,55^. These charges were combined with existing parameters for deoxyribose and phosphate groups of dA nucleotide in the Amber-OL15 force field^54^. (see Supporting Information for details).

Each system was simulated for 200 ns, during which the structures remained stable, including those containing modified nucleotides (Figure S10). To estimate chemical shifts, 50 random frames were extracted between 2 and 200 ns, and a 3.3 Å radius around T9, which includes the base-pairing partner A16 as well as nucleotides from the base-pairs immediately above and below, was used for fragment generation with AFNMR^56^ followed by GIAO-DFT calculation with Orca^57,58^. The relative chemical shift (Δδ, ppm) between HG and WCF in A_2_ wt was reliably estimated for non-exchangeable protons (A16 ^C^H8, A16 ^C^H2, and A17 ^C^H2). However, as expected, for exchangeable imino protons (T9 ^N^H3, and T8 ^N^H3), higher variability was observed (Figure S11a). Nevertheless, the mean Δδ trends were consistent with the experimental observations. In A_2_-P16, similar trends in ^1^H chemical shifts were observed, except for P16 ^C^H6, where the predicted Δδ differed in sign from the experimental value (Figure S11b). For A_2_-c^7^A16, the predicted Δδ for T9 ^N^H3 matched the experimental trend (Figure S11c), supporting the hypothesis that the minor conformation observed in *R*_1ρ_ experiments (Figure 4d, S11b) represents an HG-like state.

To further explore the structural basis of the novel ES2 implicated in the WCF-HG exchange, enhanced sampling via well-tempered parallel biased metadynamics^37–39,59,60^ was employed. This method enables the exploration of transient intermediates along the transition pathway^61–64^. Previous simulations identified two primary mechanisms for the purine *anti-*to*-syn* flip: an intrahelical pathway with a small base opening angle and an extrahelical route with a larger opening angle^3,64^. However, these simulations have primarily focused on A_6_-DNA with six contiguous A-T base-pairs and assumed a two-state exchange model with no experimental indication of an additional excited state.

To investigate ES2 in A_2_ DNA (Figure 1a and Figure 3), the orientation of the A16 nucleobase was steered using five collective variables (CVs) (Table S10, for a detailed description, see supporting information). The reweighted 2D free energy surfaces (FES) revealed two expected minima corresponding to WCF and HG conformations, as well as two additional minima potentially corresponding to ES2 (Figure S12). Structural clustering based on energy-weighted trajectories yielded representative conformers, from which chemical shifts were calculated (Figure S11d). Agglomerative clustering^65^ based on chemical shifts of all H, C and N atoms in T8, T9, G10, C15, A16 and A17 identified four major clusters (Figure 5c and Figure S12). Two chemical shift clusters mapped precisely onto the known WCF and HG conformations, while two intermediate states, Model 1 and Model 2, were located along the WCF-HG transition pathway.

Calculated Δδ values for WCF/HG, WCF/model 1, and WCF/model 2 were of similar magnitude to experimental data for non-exchangeable A16 ^C^H8 and A16 ^C^H2 protons (Figure 5d). This supports the possibility that both model 1 and model 2 represent conformers of ES2. In model 1, A16 adopts a conformation where the amino group forms a hydrogen bond with T9 O2, while in model 2, A16 is partially displaced from the helical axis (Figure 5e). In A_2_-P16, where the amino group of A16 is absent-increased exchange rate of WCF - ES2 was observed (Figure 4d). Additionally, model 2 is observed to be consistently in a high-energy region of the 2D FES surfaces for different CVs (Figure S12). These observations suggest that disruption of the A16-T9 hydrogen bond between A16 ^N^H6 and T9 O4/2, as in model 1, may be the rate-limiting step, and is the more likely structural representation of ES2.

## Conclusion

This study provides a detailed theoretical description of the influence of dipolar-coupled protons ^1^H_dip_ in proximity to protons undergoing exchange during a ^1^H *R*_1ρ_ RD experiment. We demonstrate that for ^1^H_dip_ distances greater than 3 Å, as are the most common in typical DNA and RNA structures, with Δω_dip_ either within or outside the screened offset range in an off-resonance experiment, the impact of cross-relaxation on *R*_1ρ_ is negligible. This validates the use of standard exchange models to fit the relaxation dispersion data. Several effects were studied for cases when ^1^H_dip_ is present at distances less than 3 Å and Δω_dip_ falls within the screened offset range. These findings broaden the applicability of ^1^H *R*_1ρ_ relaxation dispersion experiment, increasing the number of sites that can be probed for dynamics in nucleic acids.

Using A_2_ DNA, we detected a previously unreported second excited state (ES2) in the WCF–HG transition. MD and metadynamics simulations were applied to reveal that ES2 likely corresponds to an intermediate state featuring a hydrogen bond between the A16 amino group and T9 O2 during the WCF-HG exchange pathway. Furthermore, the increased exchange rate to ES2 observed in nebularine-modified A_2_-P16 DNA, lacking the A16 amino group, supports a mechanistic role for this interaction in modulating the transition. Detecting the previously hidden ES2 and HG conformers in both A_2_ P16- and A_2_ c^7^A16-modified DNA, highlights the improved sensitivity of ^1^H *R*_1ρ_ experiments compared to ^13^C and ^15^N *R*_1ρ_ techniques. The ES2 conformation was observed to be stabilized when A_2_ DNA was combined with Actinomycin D, suggesting its potential role in binding anticancer drugs.

The ability to characterize transient, additional states like ES2 in the WCF-HG dynamics through integrated experimental and computational approaches opens new avenues for studying conformational landscapes of nucleic acids and their roles in molecular recognition.

## Acknowledgements

R.D. acknowledges funding from European Union’s Horizon 2020 research and innovation programme under the Marie Skłodowska-Curie Action (MSCA) [101067627, project: ECONOMICS]; C. S. acknowledges funding by an EMBO postdoctoral fellowship (ALTF 1011-2020); K.P. acknowledges funding from Wallenberg Academy Fellow (KAW 2019, 0227), project grant from the Knut och Alice Wallenberg foundation (KAW 2016.0087), Cancerfonden (CAN 2018/715 & 21 1770 Pj-BF 1), KI consolidator grant (2-2111/2019) and Karolinska Institute for the help with the purchase of our 600 MHz NMR.

## Data availability

All data needed to evaluate the conclusion are present in the main text, supporting information, and supporting Excel file. Unprocessed data and code for NMR simulations can be obtained from Zenodo (10.5281/zenodo.17155221) and GitHub (https://github.com/PetzoldLab/2025_A2-Hoogsteen_public) respectively.

## Notes

https://doi.org/10.5281/zenodo.17155221

https://github.com/PetzoldLab/2025_A2-Hoogsteen_public

### Competing Interest Statement

The authors have declared no competing interest.

