## Supporting information for "^1^H *R*_1ρ_ Relaxation Identifies a Hidden Intermediate in DNA Base-Pairing"

[§] Equal contribution

### *Material and Methods*

#### *Solid phase synthesis of modified DNA*

5'-*O*-DMT-protected 3'-β-cyanoethyl phosphoramidites of dA, dC, dG, dT, dP (2'-deoxynebularine) and $c^7$dA, as well as the 5'-*O*-MMT-protected 3'-β-cyanoethyl $m^1$dA phosphoramidite, were purchased from Glen Research, while other reagents were obtained from Sigma Aldrich.

Solid-phase synthesis of DNA oligonucleotides was carried out using standard phosphoramidite chemistry on a 1 μmol scale with 5-(ethylthio)-1*H*-tetrazole (ETT) as the activator. Phosphoramidites were employed as 50 or 70 mM (standard or modified DNA nucleotides, respectively) solutions in anhydrous MeCN.

Cleavage from the solid support and removal of base-labile protecting groups was achieved by treatment with a 1:1 mixture of 25% aq. $NH_3$ and 40% aq. $MeNH_2$ (1.0 mL) at ambient temperature for 30 min, after which the support was washed with an additional 0.5 mL of the same mixture. The combined solutions were heated to 65 °C for 30 min.

For $m^1$dA-modified DNA, $N^6$-phenoxyacetyl and $N^2$-(4-isopropyl)phenoxyacetyl protecting groups were employed on the dA and dG phosphoramidites, respectively. Capping was performed with a combination of 5% phenoxyacetic anhydride in THF/pyridine and 16% 1-methylimidazole in THF and 20 mM $I_2$ in THF/$H_2O$/pyridine was used as the oxidizer[1]. After drying under reduced pressure, the solid support was treated with 2 M $NH_3$ in MeOH (1.0 mL) at ambient temperature for 24 h. It was washed with an additional 0.5 mL of the same solution; the combined solutions were evaporated to dryness and the residue was dissolved in 1.5 mL of $H_2O$.

The crude solid-phase synthesis products were purified using Glen-Pak DNA purification cartridges according to the manufacturer's instructions. $m^1$dA-modified DNA was eluted from the cartridge with 0.5% $NEt_3$ in MeCN/$H_2O$. Oligonucleotides were analysed by 20% denaturing PAGE. Yields were determined by UV absorbance and are reported in Table S2.

#### *NMR sample preparation*

Both strands of the unmodified $A_2$ DNA ($A_2$-wt) duplex were purchased from Integrated DNA Technology (IDT) as standard desalting. The strands were mixed to a final concentration of 1.5 mM in 500 μL NMR buffer (15 mM $NaP_i$ pH 6.5, 25 mM NaCl, 0.1 mM EDTA). This solution was subjected to slow annealing during which it was heated to 95 °C for 5 min, followed by incubation at 65 °C, 37 °C, 25 °C and finally 4 °C for 30 min each. For modified $A_2$ DNA ($A_2$-P16, $A_2$-$c^7$A16 and $A_2$-$m^1$A16), the purified samples were buffer-exchanged to NMR buffer after annealing. The final samples were concentrated to the respective concentrations reported in Table S3 in a final volume of 250 μL.

Actinomycin D (ActD) was purchased from Sigma Aldrich (A9415-2MG) and used without any further purification. 2 mg of powder was dissolved in 160 μL of NMR buffer + 840 μL of MeOH giving a stock solution of 1.6 mM. 156 μL of this solution was evaporated under reduced pressure and the remaining solid was resuspended in 250 μL of 1 mM unmodified $A_2$ DNA in NMR buffer giving a 1:1 $A_2$ DNA:ActD solution. This solution was spiked with 6% $D_2O$ and transfered to in a 5 mm Shigemi tube for NMR measurements.

***NMR measurements and processing***

NMR measurements were performed on a 600.16 MHz Avance III-HD Bruker spectrometer equipped with a QCI-P cryoprobe. SOFAST HMQC[2,3] spectra were acquired with 128 increments and recorded with 64 scans each. The carrier frequencies of $^{1}$H and $^{13}$C were placed at 7.8 ppm and 142 ppm, respectively. Imino NOESY experiments were performed using the pulse sequence implemented in NMRlib[4]. The $^{1}$H carrier frequency was set to 13 ppm, and the excitation bandwidth was 4 ppm. $^{1}$H $R_{1\rho}$ RD experiments were performed using previously reported pulse sequences[5,6]. The acquisition parameters are tabulated in Table S6. NMR spectra were processed using Bruker Topspin 3.6.3. Deconvoluted peak intensities obtained using “mdcon” and the corresponding signal-to-noise ratios calculated with “sino” in Topspin were exported to plain text for further data analysis. Temperature-dependent $R_{1\rho}$ experiments were performed in steps of 5 K from 283 to 303 K for wild type $A_2$ DNA. $^{1}$H $R_{1\rho}$ RD experiments on modified $A_2$ DNA duplexes were performed at 278 K.

***$R_{1\rho}$ Data fitting***

Peak intensities ($I$) as a function of spin-lock duration $\tau_{SL}$ were fitted to a mono-exponential decay to obtain the underlying rotating frame relaxation rates $R_{1\rho}$. Data points were weighted according to their uncertainties given by the root mean square of the baseline noise of each spectrum, which is defined as rms = $I$ / (2 * sino). Confidence intervals for the resulting $R_{1\rho}$ values were estimated from the standard deviation obtained by Monte Carlo resampling of the original datasets with 500 replicas.

For analysis of conformational exchange, $R_{1\rho}$ was fitted as a function of spin-lock strength $\omega_{SL}$ and offset $\Omega_{SL}$ with weights given by the standard deviation. Exchange processes were modelled as two-state or three-state without or with minor state exchange by taking $R_{1\rho}$ as the least negative real eigenvalue of the corresponding Bloch-McConnel matrix (see below for a full description of the matrices)[7]. Models were ranked using the Akaike information criterion with small sample size correction (AICc), Bayesian information criterion (BIC, difference $\geq 10$) and $F$-test (95% confidence level)[8]. The goodness of fit was further judged based on the distribution of fit residuals, which are ideally random and centred around $y = 0$ for a model that captures the exchange process appropriately. Confidence intervals for each exchange parameter of the best model were estimated from the respective standard deviation obtained by Monte Carlo resampling of the original datasets with 500 replicas.

All data analysis was performed using a custom program written in Python 3.10. Linear algebra operations for obtaining exact numerical eigenvalues were implemented with the NumPy package. Non-linear least-squares minimization with the Levenberg-Marquardt algorithm was performed using the lmfit package[9]. The attached excel sheets provides the data for the individual and global fits for the protons subjected to $^{1}$H $R_{1\rho}$ experiments. The AICc and BIC criteria are reported for each of the three-state models. The best model from these statistical tests and the F-test is highlighted with thick border.

***Molecular Dynamics (MD) simulation***

MD simulations of $A_2$ DNA were performed for both the WCF and HG conformations of the nucleobase A16. The lowest energy structure from PDB id: 5UZD[10] was used as a model for the WCF conformation. For the HG conformation the glycosidic $\chi$ dihedral angle of A16 was rotated by 180°. The Amber14SB force field with OL15 parameters for DNA (https://fch.upol.cz/ff_ol/gromacs.php) was used to simulate both the models using the GROMACS 2023.4 suite. All-atom MD simulations were performed for 200 ns using a TIP3P water model in a dodecahedral box. The system was first neutralized using of $Na^+$ ions and the final NaCl concentration was set to 25 mM. The system was energy minimized via the steepest decent gradient method to a maximum force constant < 1000 kJ/mol, equilibrated for a total of 200 ps at 300 K using a velocity rescale scheme and 1 bar pressure using the Parrinello-Rahman barostat. A Verlet nonbonded cut-off scheme with grid neighbour search and 1.0 nm cut-off for van der Waals interaction with energy and pressure dispersion correction was used. Particle Mesh Ewald with fourth order cubic interpolation was used for Coulomb interactions. All bonds were constrained using the LINCS algorithm. During the MD run, a 2fs integration step was used and the trajectory was saved every 10 ps. The trajectory was analysed using Plumed 2.7[11,12], UCSF Chimera 1.17.3 and Python 3.11. Simulations were performed in the Tetralith HPC cluster via the National Academic Infrastructure for supercomputing in Sweden (NAISS). PDB models and molecular dynamics parameters files are provided in Zenodo (10.5281/zenodo.17155221).

***Parameterization of modified nucleotides for MD simulation***

To derive force field parameters for deoxyribonucleotides of purine (P, residue code PRN) and 7-deazaadenine ($c^7$A, residue code 7DA), geometries of the respective $N^9$-methylated nucleobases were first optimized in the gas phase with tight convergence criteria and characterized as stationary points on the potential energy surface by analytical vibrational frequency calculation using Orca 5.0.4[13,14]. The nucleobases were subjected to two-stage restrained electrostatic potential (RESP) fitting with default parameters and standard hyperbolic restraints[15,16] using Psi4 1.9.1[17,18]. For this, the charge of the capping exocyclic methyl group was set to +0.1053 and 20 equally weighted, randomized orientations of the nucleobases were considered. GROMACS-compatible .itp files were generated with ACPYPE[19] (options -c user -a amber -o gmx) and combined with deoxyribose and phosphate parameters for unmodified adenine deoxyribonucleotide (residue code DA) as defined in the Amber force field.

***Metadynamics simulation***

GROMACS 2021.3 patched with PLUMED 2.7.2 was used to perform a parallel biased well-tempered metadynamics simulation by biasing five collective variables (Table S11). The simulation was performed for 200 ns until convergence with the height of the gaussians set to 0.5 kJ/mol deposited every 500 steps. The sigma of the gaussian height was set to the ½ of the standard deviation obtained from the unbiased simulations. The 1D free-energy surface (FES) for each collective variable (CV) were obtained from the reweighted bias energy while the 2D FES were obtained by using binned_statistic_2d from scipy.stats module of scipy using each combination of CV and the reweighted bias energy (Figure S12). All the files used for the simulations and analysis are provided in the GitHub repository (link).

***Chemical shift calculation***

Calculation of chemical shifts for T8, T9, G10, C15, A16 and A17 from MD frames was done with the help of an automated fragment generation approach[20]. Before compiling version 1.8 of the AFNMR program, the value of 'MAXNRES' in afnmr-F90 was updated to include the modified nucleotides, and their residue codes were added in the form of 'nresn(34) = 'PRN'' etc. The compiled program was checked using the built-in test suite. AFNMR requires Amber-compatible frcmod and .lib

files for modified nucleotides, which were generated using ACPYPE (options *-c user -a amber -o gmx*) from *.mol2* files of the respective residues with charges as derived during MD parametrization.
50 frames from the unbiased simulation run were selected randomly for the chemical shift calculations. The initial 2 ns trajectory were discarded due to significant change in the backbone RMSD (Figure S10). Frames from the metadynamics runs were obtained by clustering the trajectory using gromos method in GROMACS with the backbone cutoff of 0.2 nm. The output clusters were then weighted according to the reweighted energy from the metadynamics run. The central frame from each of the weighted cluster consisting of ≥ 3 frames was used for the chemical shift calculations.

AFNMR was run on each frame with the following options:
*-list "{{8..10},{15..17}" -orca -nomin -mixedb -frcmod ${MOD_NAME}_AC.frcmod -offlib ${MOD_NAME}_AC.lib -workdir.*
This produces residue-centric fragments that include all neighbouring residues, water molecules and ions in a sphere of 3.3 Å, i.e. the direct base pairing partner plus residues from the base pair above and below. The rest of the system is projected onto the fragment as a set of point charges. Chemical shielding tensors were calculated in Orca 5.0.4[13,14] by GIAO-DFT with the OLYP functional using a pcSseg-2 basis set on the central residue, pcSseg-1 basis set on all other atoms[21] and def2/J auxiliary basis set[22] for the RI-J approximation. Chemical shifts were obtained using the built-in referencing routine of AFNMR but were not corrected further as only relative chemical shift changes between MD frames/conformations were of interest.

For analysis of chemical shift changes from conventional MD trajectories of $A_2$-wt, $A_2$-P16, and $A_2$-c$^7$A16 with nucleotide 16 either in the WCF or in the HG conformation, chemical shifts of each atom type were averaged over all 50 frames and standard deviations were calculated.

For metadynamics trajectories, only output frames stemming from at least three clustered input frames were considered in the chemical shift analysis. The relevant frames were subjected to agglomerative clustering under a Euclidean distance metric with Ward linkage. Clustering was performed on all $^1$H, $^{13}$C and $^{15}$N chemical shifts of the six central residues. Different numbers of output clusters were tested and assessed by how well geometric clusters (WCF, HG and other conformations) are recovered from the chemical shift data; best results were found with four output clusters. When mapping these back onto the five CVs from metadynamics, distinct conformers for WCF and HG base pairing as well as two additional arrangements (Model 1, Model 2) are observed.

#### *Effects of cross-relaxations on the $R_{1\rho}$ rates without chemical exchange*

The Bloch equation to model the magnetization behavior under a spin-lock for a single spin in the rotating frame is given by[8]

$$\frac{\mathrm{d}}{dt}\begin{bmatrix}\mathrm{M}_x(t)\\ \mathrm{M}_y(t)\\ \mathrm{M}_z(t)\end{bmatrix} = -1\begin{bmatrix}R_2 & \Omega & 0\\ -\Omega & R_2 & \omega_{\mathrm{SL}}\\ 0 & -\omega_{\mathrm{SL}} & R_1\end{bmatrix}\begin{bmatrix}\mathrm{M}_x(0)\\ \mathrm{M}_y(0)\\ \mathrm{M}_z(0)\end{bmatrix} \tag{S1}$$

Where $M_x$, $M_y$ and $M_z$ represents the bulk magnetization along the x, y and z axis, $R_2$ and $R_1$ are the transversal and longitudinal auto-relaxation rates respectively. The off-diagonal terms describe the precession frequency Ω along the z-axis and spin-lock field $\omega_{SL}$ along the x-axis. Rotating equation S1 to the spin-locking field or the double rotating frame around y-axis by an angle θ with respect to z-axis, provides the rate equation for the magnetization along each axis.

$$\frac{\mathrm{d}}{\mathrm{d}t}\begin{bmatrix} \mathrm{M}'_x(t) \\ \mathrm{M}'_y(t) \\ \mathrm{M}'_z(t) \end{bmatrix} = -1\begin{bmatrix} R_1\sin^2\theta + R_2\cos^2\theta & \sqrt{\Omega^2+\omega^2} & (R_2-R_1)\sin\theta\cos\theta \\ -\sqrt{\Omega^2+\omega^2} & R_2 & 0 \\ (R_2-R_1)\sin\theta\cos\theta & 0 & R_1\cos^2\theta + R_2\sin^2\theta \end{bmatrix}\begin{bmatrix} \mathrm{M}'_x(0) \\ \mathrm{M}'_y(0) \\ \mathrm{M}'_z(0) \end{bmatrix} \quad \text{(S2)}$$

Under the experimentally condition of $\omega_{SL} >> (R_2 - R_1)\sin\theta\cos\theta$ and due to rapid inter-conversion of Mx' and My' because of $\sqrt{\Omega^2+\omega^2}$, the off-diagonal term $(R_2 - R_1)\sin\theta\cos\theta$ becomes 0. This provides the rate equation for the Mz' under the spin-locking field

$$\frac{\mathrm{d}}{\mathrm{d}t}[\mathrm{M}'_z(t)] = -R_{1\rho}\mathrm{M}'_z(0) \quad \text{(S3)}$$

The general solution to equation S3 is

$$\mathrm{M}'_z(t) = \exp(-R_{1\rho}t)\mathrm{M}'_z(0) \quad \text{(S4)}$$

here, $t$ is the duration of the spin-locking field which can be represented as $\tau_{SL}$ and $R_{1\rho}$ is given by

$$R_{1\rho} = R_1\cos^2\theta + R_2\sin^2\theta \quad \text{(S5)}$$

This equation shows that the Mz' magnetization will decay monoexponentially with the rate of $R_{1\rho}$ which is then given as the weighted sum of the $R_1$ and $R_2$ rates. To ascertain the effect of longitudinal and transverse cross-relaxation rates (σ and μ) between two spins (a and b), the matrix in equation S1 is expanded to

$$\frac{\mathrm{d}}{\mathrm{d}t}\begin{bmatrix} \mathrm{Ma}_x(t) \\ \mathrm{Ma}_y(t) \\ \mathrm{Ma}_z(t) \\ \mathrm{Mb}_x(t) \\ \mathrm{Mb}_y(t) \\ \mathrm{Mb}_z(t) \end{bmatrix} = -1\begin{bmatrix} R_{2a} & \Omega & 0 & \mu & 0 & 0 \\ -\Omega & R_{2a} & \omega_{SL} & 0 & \mu & 0 \\ 0 & -\omega_{SL} & R_{1a} & 0 & 0 & \sigma \\ \mu & 0 & 0 & R_{2b} & \Omega & 0 \\ 0 & \mu & 0 & -\Omega & R_{2b} & \omega_{SL} \\ 0 & 0 & \sigma & 0 & -\omega_{SL} & R_{2b} \end{bmatrix}\begin{bmatrix} \mathrm{Ma}_x(0) \\ \mathrm{Ma}_y(0) \\ \mathrm{Ma}_z(0) \\ \mathrm{Mb}_x(0) \\ \mathrm{Mb}_y(0) \\ \mathrm{Mb}_z(0) \end{bmatrix} \quad \text{(S6)}$$

Here,

$$\mu = \mathrm{D}\,(2\mathrm{J}(0) + 3\mathrm{J}(\omega)) \quad \text{and} \quad \sigma = \mathrm{D}\,(-\mathrm{J}(0) + 6\mathrm{J}(2\omega))$$

$$\mathrm{D} = \frac{1}{4}\left(\frac{\gamma_H^2\hbar\mu_0}{4\pi}\right)^2\frac{1}{r^6} \quad \text{and} \quad \mathrm{J}(\omega) = \frac{2}{5}\frac{\tau_c}{(1+\omega^2\tau_c^2)} \quad \text{(S7)}$$

Where $\tau_c$ is the rotation correlation time, ω is the proton Larmor frequency, $\gamma_H$ is the proton gyromagnetic ratio, $\hbar$ is the reduced Planck's constant, $\mu_0$ is the vacuum permeability, and $r$ is the inter-proton distance. Rotating the matrix in equation S6 around y-axis into the double rotating frame, the $R_{1\rho}$ is given as

$$R_{1\rho} = (R_1 + \sigma)\cos^2\theta + (R_2 + \mu)\sin^2\theta \quad \text{(S8)}$$

This suggests that the cross-relaxation rates will scale the $R_1$ and $R_2$ contribution to the $R_{1\rho}$ rate but do not affect the mono-exponential behaviour of the z-magnetization decay under spin-locking field. Equation S8 can be now expanded to include chemical exchange as shown below.

***Bloch-McConnel (BM) matrix for two-state exchange***

The BM matrix for a two-state chemical ES (b) ⇌ GS (a) exchange is given by[8]

$$\frac{\mathrm{d}}{\mathrm{d}t}\begin{bmatrix} \mathbf{M}_a(t) \\ \mathbf{M}_b(t) \end{bmatrix} = \left\{\begin{bmatrix} \mathbf{L}_a & \mathbf{0} \\ \mathbf{0} & \mathbf{L}_b \end{bmatrix} + \mathbf{K}\otimes\mathbf{1}\right\}\begin{bmatrix} \mathbf{M}_a(0) \\ \mathbf{M}_b(0) \end{bmatrix} \quad \text{(S9)}$$

$$\mathbf{L}_{i(a,b)} = -1\begin{bmatrix} R_{2i} & \delta_i & -\omega_{\text{ramp}} \\ -\delta_i & R_{2i} & \omega_{\text{SL}} \\ \omega_{\text{ramp}} & -\omega_{\text{SL}} & R_{1i} \end{bmatrix} \qquad \begin{aligned} \delta_{\text{a}} &= -p_{\text{b}}\,\Delta\omega_{\text{b}} - \Omega_{\text{SL}} \\ \delta_{\text{b}} &= p_{\text{a}}\,\Delta\omega_{\text{b}} - \Omega_{\text{SL}} \end{aligned} \tag{S10}$$

$$\mathbf{K} = \begin{bmatrix} -k_{12} & k_{21} \\ k_{12} & -k_{21} \end{bmatrix} \qquad \begin{aligned} k_{12} &= p_{\text{b}} k_{\text{ex}} \\ k_{21} &= p_{\text{a}} k_{\text{ex}} \end{aligned} \tag{S11}$$

$$[\mathbf{M}_{\text{a}}(0) \quad \mathbf{M}_{\text{b}}(0)]^{\text{T}} = \left[\text{M}_{x(\text{a})} \quad \text{M}_{y(\text{a})} \quad \text{M}_{z(\text{a})} \quad \text{M}_{x(\text{b})} \quad \text{M}_{y(\text{b})} \quad \text{M}_{z(\text{b})}\right]^{\text{T}} \tag{S12}$$

Where $R_2$ is the transverse relaxation rate, $R_1$ is the longitudinal relaxation rate, $p_a$ is the population of the GS, $p_b$ is the population of the ES, $k_{ex}$ ($k_{ab} + k_{ba}$) is the exchange rate, $\Delta\omega_b$ is the offset of the ES with respect to the observed signal, $\Omega_{SL}$ is the offset of the spin-lock with respect to the observed signal, $\omega_{SL}$ is the spin-lock strength and $\omega_{ramp}$ is the strength of the ramp pulse flanking the spin-lock. The superscript $^{T}$ denotes the transpose of the initial magnetization vector. **0** and **1** are the 3 x 3 zero and identity matrix, respectively. The solution to equation S9 is given by,

$$\begin{bmatrix} \mathbf{M}_{\text{a}}(t) \\ \mathbf{M}_{\text{b}}(t) \end{bmatrix} = \exp\left(\left\{\begin{bmatrix} \mathbf{L}_{\text{a}} & \mathbf{0} \\ \mathbf{0} & \mathbf{L}_{\text{b}} \end{bmatrix} + \mathbf{K} \otimes \mathbf{1}\right\} * \tau_{\text{SL}}\right) \cdot \begin{bmatrix} \mathbf{M}_{\text{a}}(0) \\ \mathbf{M}_{\text{b}}(0) \end{bmatrix} \tag{S13}$$

Where $\tau_{SL}$ is the duration of the spinlock. Assuming that, $R_{1a} = R_{1b}$ and $R_{2a} = R_{2b}$, $R_{1\rho}$ is estimated by fitting a mono-exponential decay to

$$M_{z(a)}(t) = M_{z(a)}(0)\exp\left(-R_{1\rho} * \tau_{\text{SL}}\right) \tag{S14}$$

with, $M_{z(a)}(0) = p_a$. The analytical approximation of the BM matrix to describe $R_{1\rho}$ was derived to be[8,23]:

$$R_{1\rho} = R_1\cos^2\theta + R_2\sin^2\theta + R_{ex}\sin^2\theta \tag{S15}$$

Here, $\theta = \arctan\left(\frac{\omega_{\text{SL}}}{\Omega_{\text{SL}}}\right)$. Rearranging equation S15 gives the expression for the $R_{2\text{eff}}$ or $R_2 + R_{\text{ex}}$,

$$R_{2\text{eff}} = R_2 + R_{\text{ex}} = \frac{R_{1\rho} - R_1\cos^2\theta}{\sin^2\theta} \tag{S16}$$

***Extended two-state Bloch-McConnell matrix***

Equation S9 can be extended to include cross-relaxation, and can be written as[24]:

$$\frac{\text{d}}{\text{d}t}\begin{bmatrix} \mathbf{M}_{\text{a}}(t) \\ \mathbf{M}_{\text{b}}(t) \\ \mathbf{M}_{\text{c}}(t) \end{bmatrix} = \left\{\begin{bmatrix} \mathbf{L}_{\text{a}} & \mathbf{0} & \mathbf{L}_{\text{CR}} \\ \mathbf{0} & \mathbf{L}_{\text{b}} & \mathbf{L}_{\text{CR}} \\ \mathbf{L}_{\text{CR}} & \mathbf{L}_{\text{CR}} & \mathbf{L}_{\text{c}} \end{bmatrix} + \mathbf{K} \otimes \mathbf{1}\right\}\begin{bmatrix} \mathbf{M}_{\text{a}}(0) \\ \mathbf{M}_{\text{b}}(0) \\ \mathbf{M}_{\text{c}}(0) \end{bmatrix} \tag{S17}$$

Here, $\mathbf{M_c}$ is the magnetization vector for $^1H_{dip}$ with the cross-relaxation given by matrix $\mathbf{L_{CR}}$.

$$\mathbf{L}_{\text{CR}} = -1\begin{bmatrix} \mu & 0 & 0 \\ 0 & \mu & 0 \\ 0 & 0 & \sigma \end{bmatrix} \tag{S18}$$

The distance from GS and ES to the dipolar proton can be modulated using equation S17 and S18. The $R_{1\rho}$ is given by

$$R_{1\rho} = (R_1 + \sigma)\cos^2\theta + (R_2 + \mu)\sin^2\theta + R_{ex}\sin^2\theta \tag{S19}$$

This, again, describes a mono-exponential decay of spin-locked magnetization in the presence of cross-relaxation.

***Three-state exchange model without cross-relaxation***

The three-state exchange with was modelled using the following Bloch-McConnell matrix[8]:

$$\frac{\mathrm{d}}{\mathrm{d}t}\begin{bmatrix}\mathbf{M}_a(t)\\ \mathbf{M}_b(t)\\ \mathbf{M}_c(t)\end{bmatrix} = \left\{\begin{bmatrix}\mathbf{L}_a & \mathbf{0} & \mathbf{0}\\ \mathbf{0} & \mathbf{L}_b & \mathbf{0}\\ \mathbf{0} & \mathbf{0} & \mathbf{L}_c\end{bmatrix} + \mathbf{K}\otimes\mathbf{1}\right\}\begin{bmatrix}\mathbf{M}_a(0)\\ \mathbf{M}_b(0)\\ \mathbf{M}_c(0)\end{bmatrix} \quad \text{(S20)}$$

$$\mathbf{L}_{\mathrm{i(a,b,c)}} = -1\begin{bmatrix} R_{2i} & \Delta_i & -\omega_{ramp}\\ -\Delta_i & R_{2i} & \omega_{\mathrm{SL}}\\ \omega_{ramp} & -\omega_{\mathrm{SL}} & R_{1i}\end{bmatrix} \qquad \begin{aligned}\Delta_a &= -\,p_b\Delta\omega_b - p_c\Delta\omega_c - \Omega_{\mathrm{SL}}\\ \Delta_b &= (1 - p_b\Delta\omega_b) - p_c\Delta\omega_c - \Omega_{\mathrm{SL}}\\ \Delta_c &= (1-p_c)\Delta\omega_c - p_b\Delta\omega_b - \Omega_{\mathrm{SL}}\end{aligned} \quad \text{(S21)}$$

Different topologies of the three-state exchange can be modelled by changing the rate matrix **K**. For a general triangular topology (ES1 (b) ⇌ GS (a) ⇌ ES2 (c) ⇌ ES1 (b)), the K matrix is given by:

$$\mathbf{K} = \begin{bmatrix} -k_{12} - k_{13} & k_{21} & k_{31}\\ k_{12} & -k_{21} - k_{23} & k_{32}\\ k_{13} & k_{23} & -k_{31} - k_{32}\end{bmatrix} \quad \text{(S22)}$$

$$\begin{aligned} k_{12} &= k_{ex\_ab}\frac{p_b}{p_a + p_b} & k_{21} &= k_{ex\_ab}\frac{p_a}{p_a + p_b}\\ k_{13} &= k_{ex\_ac}\frac{p_c}{p_a + p_c} & k_{31} &= k_{ex\_ac}\frac{p_a}{p_a + p_c}\\ k_{23} &= k_{ex\_bc}\frac{p_c}{p_b + p_c} & k_{32} &= k_{ex\_bc}\frac{p_b}{p_b + p_c}\end{aligned} \quad \text{(S23)}$$

For a three-state linear topology (GS (a) ⇌ ES1 (b) ⇌ ES2 (c)), $k_{31}$ and $k_{13}$ = 0, while for the star-like topology (ES1 (b) ⇌ GS (a) ⇌ ES2 (c)), $k_{32}$ and $k_{23}$ = 0.

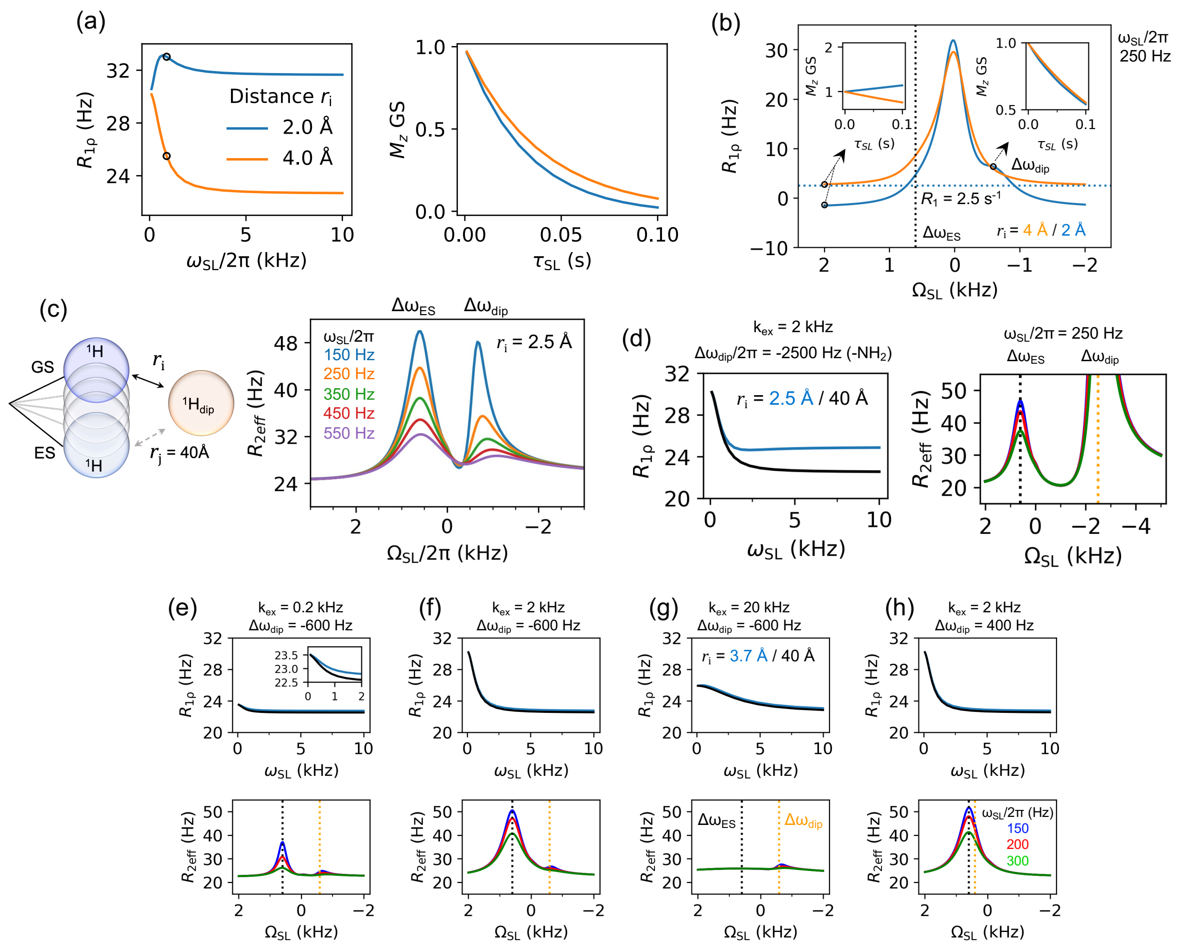


**Figure S1**. (a) Representative on-resonance $R_{1\rho}$ curves (depicting an initial rise for a $^1H_{dip}$ at $r_i$ = 2 Å (blue) as compared to $r_i$ = 4 Å (orange), demonstrating the extent of μ contribution for smaller distances. The associated behaviour of $M_z$ magnetization in the GS is plotted against $\tau_{SL}$ at $\omega_{SL}$ = 200 Hz (black circles, left). (b) Simulated off-resonance $R_{1\rho}$ curves for $r_j$ = 40 Å, with $r_i$ = 4 Å (orange) $r_i$ = 2 Å (blue) at $\omega_{SL}$ = 250 Hz. Inserts depict $M_{z,\,GS}$ behaviour at offsets indicated by black circles and arrows, highlighting the rise in magnetisation at $r_i$ = 2 Å. $\Delta\omega_{ES}$ is marked with a dotted black line while a horizontal blue dashed line indicates $R_1$. $\Delta\omega_{dip}$ is also denoted on the blue curve. (c) Simulated off-resonance $R_{2eff}$ ($R_2 + R_{ex}$) at different $\omega_{SL}/2\pi$ (150 Hz blue, 250 Hz orange, 350 Hz green, 450 Hz brown and 550 Hz purple), for $r_i$ = 2.5 Å and $r_j$ = 40 Å (no cross-relaxation in ES). The local maximum at $\Delta\omega_{dip}$ shifts with changing $\omega_{SL}$ while $\Delta\omega_{ES}$ remains constant. Simulation parameters: $k_{ex}$ = 2 kHz, $p_{ES}$ = 0.5%, $\tau_c$ = 5.1 ns, $R_1$ = 2.5 s$^{-1}$, $R_2$ = 22.5 s$^{-1}$, $\Delta\omega_{ES}/2\pi$ = 600 Hz, $\Delta\omega_{dip}/2\pi$ = −600 Hz. (d) Simulation of on-resonance $R_{1\rho}$ and off-resonance $R_{2eff}$ curves where $^1H_{dip}$ represents amino protons (-$NH_2$) at $r_i$ = 2.5 Å from imino protons, as canonically observed in WCF-base-paired B-form DNA or A-form RNA helices. The average $\Delta\omega_{dip}$ between the imino and amino protons is set to be −2500 Hz with $\Delta\omega_{ES}$ = +600 Hz. The inclusion of an amino proton increases $R_2$ to $R_2$ + μ (see main text), but the large $\Delta\omega_{dip}$, minimizes any significant effect on the exchange parameters between GS and ES. (e–h) Simulated on- and off-resonance profiles for $r_j$ = 40 Å and either $r_i$ = 3.7 Å (light blue, representing conventional inter-imino proton distances in B-form DNA or A-form RNA) or $r_i$ = 40 Å (black, representing no dipolar-coupled protons). Various combinations of $k_{ex}$ and $\Delta\omega_{dip}$ were simulated. $\Delta\omega_{ES}$ and $\Delta\omega_{dip}$ are indicated with dashed black and orange lines, respectively, in the off-resonance plots. For panel (h) off-resonance curves were simulated at $\omega_{SL}$ = 150 Hz (blue), 200 Hz (red) and 300 Hz (green). The effects of $^1H_{dip}$ are negligible under most exchange conditions except when $\Delta\omega_{ES}$ and $\Delta\omega_{dip}$ have the same sign, as shown in panel (h) (see main text for details).

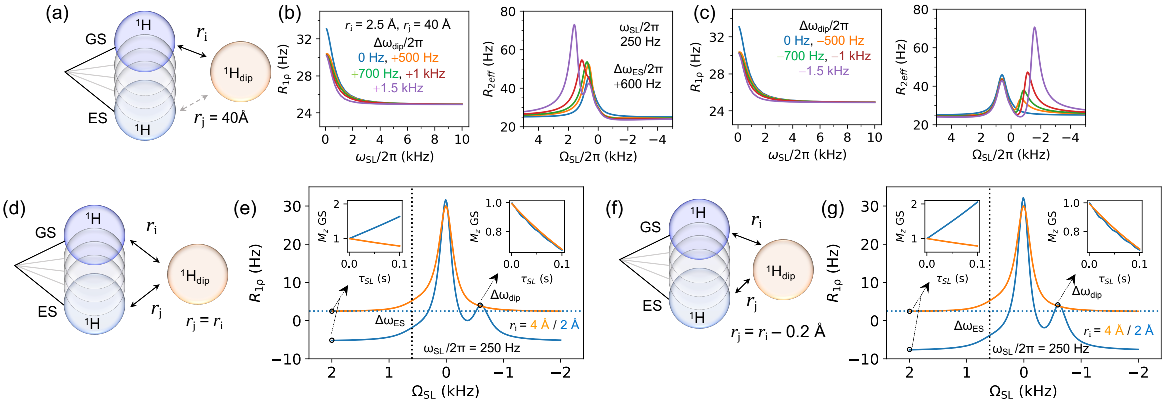


**Figure S2**. (a) Model representing scenario 1 where $r_i$ = 2.5 Å and $r_j$ = 40 Å (no cross relaxation with ES). (b, c) Simulated on- and off-resonance profiles for various $\Delta\omega_{dip}$ values (0 Hz blue, ±500 Hz orange, ±700 Hz green, ±1 kHz brown, and ±1.5 kHz purple), with $\Delta\omega_{ES}/2\pi$ = 600 Hz and $\omega_{SL}/2\pi$ = 250 Hz. These simulations illustrate the effect of $^1H_{dip}$ at different chemical shifts of $^1H_{dip}$ relative to the excited state (ES). When $\Delta\omega_{dip}$ and $\Delta\omega_{ES}$ have opposite signs (right) exchange parameters can be estimated with high confidence; however, when their signs are same (left), accurate estimation becomes more challenging. Model representing scenario 2 (d) where $r_i$ = 2.5 Å = $r_j$ and scenario 3 (f) where $r_i$ = 2.5 Å and $r_j$ = $r_i$ – 0.2 Å. (e, g) Simulated off-resonance $R_{1\rho}$ plots for varying distances with $r_i$ of 4 Å (orange) or 2 Å (blue). Inserts depict the behaviour of $M_{z,GS}$ at the offsets marked by black circles. The position of $\Delta\omega_{ES}$ is indicated with a dotted black line, while the blue dashed horizontal line marks $R_1$. All simulations were conducted with $\omega_{SL}/2\pi$ = 250 Hz using the parameters described in Figure S1. These plots reveal that strong dipolar coupling between $^1H_{dip}$ and both the GS and ES leads to an exponential rise in $R_{1\rho}$ rather than the expected exponential decay with respect to the spinlock duration ($\tau_{SL}$). At extreme offsets and short distances (e.g., 2 Å), this coupling can result in apparent negative $R_{1\rho}$ values, causing the rates to deviate significantly from $R_1$.

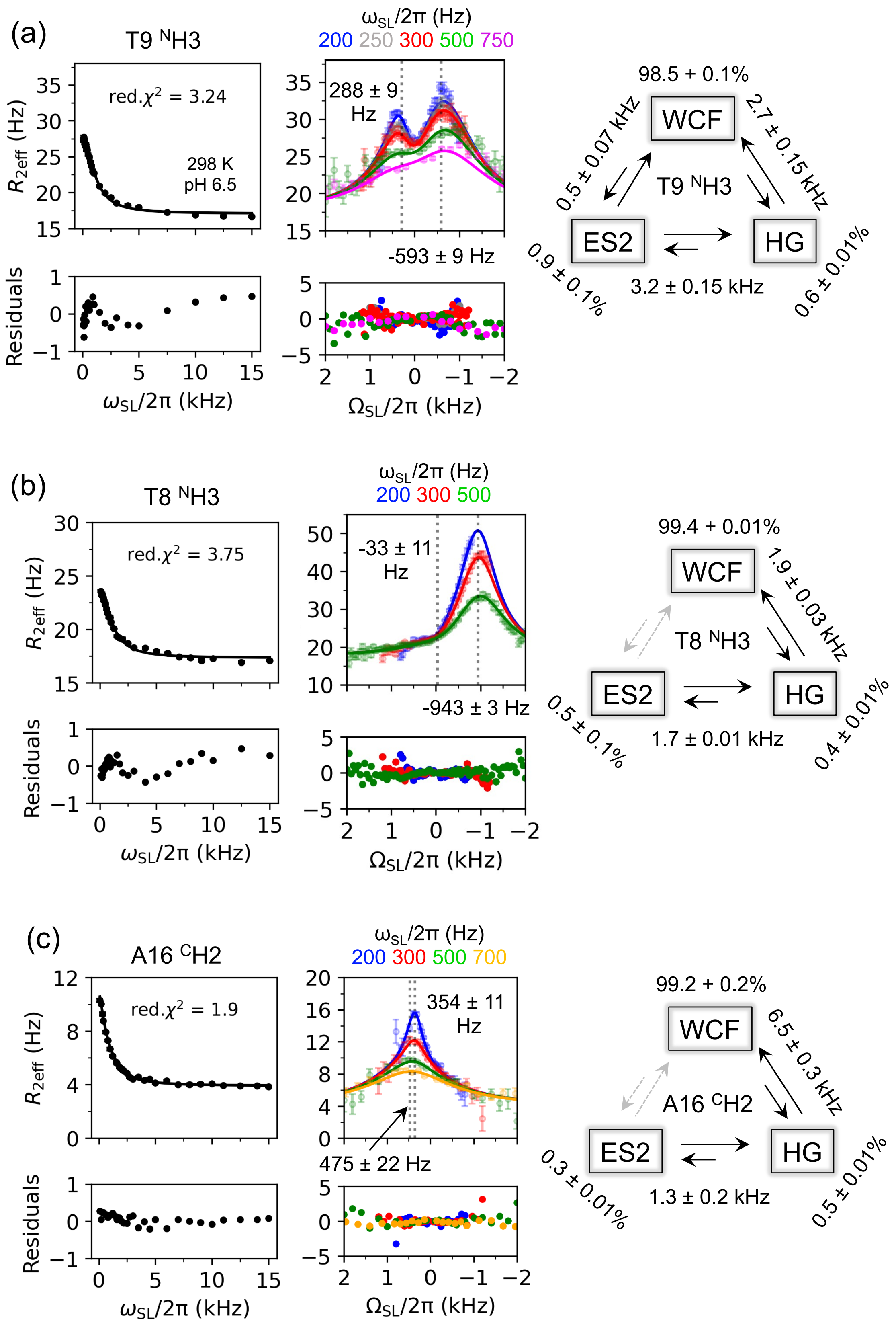

(a)
T9 NH3
red.χ2 = 3.24
298 K
pH 6.5
R2eff (Hz)
Residuals
ωSL/2π (kHz)
ωSL/2π (Hz)
200 250 300 500 750
288 ± 9 Hz
-593 ± 9 Hz
ΩSL/2π (kHz)
98.5 + 0.1%
WCF
0.5 ± 0.07 kHz
2.7 ± 0.15 kHz
T9 NH3
ES2
HG
3.2 ± 0.15 kHz
0.9 ± 0.1%
0.6 ± 0.01%
(b)
T8 NH3
red.χ2 = 3.75
R2eff (Hz)
Residuals
ωSL/2π (kHz)
ωSL/2π (Hz)
200 300 500
-33 ± 11 Hz
-943 ± 3 Hz
ΩSL/2π (kHz)
99.4 + 0.01%
WCF
1.9 ± 0.03 kHz
T8 NH3
ES2
HG
1.7 ± 0.01 kHz
0.5 ± 0.1%
0.4 ± 0.01%
(c)
A16 CH2
red.χ2 = 1.9
R2eff (Hz)
Residuals
ωSL/2π (kHz)
ωSL/2π (Hz)
200 300 500 700
354 ± 11 Hz
475 ± 22 Hz
ΩSL/2π (kHz)
99.2 + 0.2%
WCF
6.5 ± 0.3 kHz
A16 CH2
ES2
HG
1.3 ± 0.2 kHz
0.3 ± 0.01%
0.5 ± 0.01%

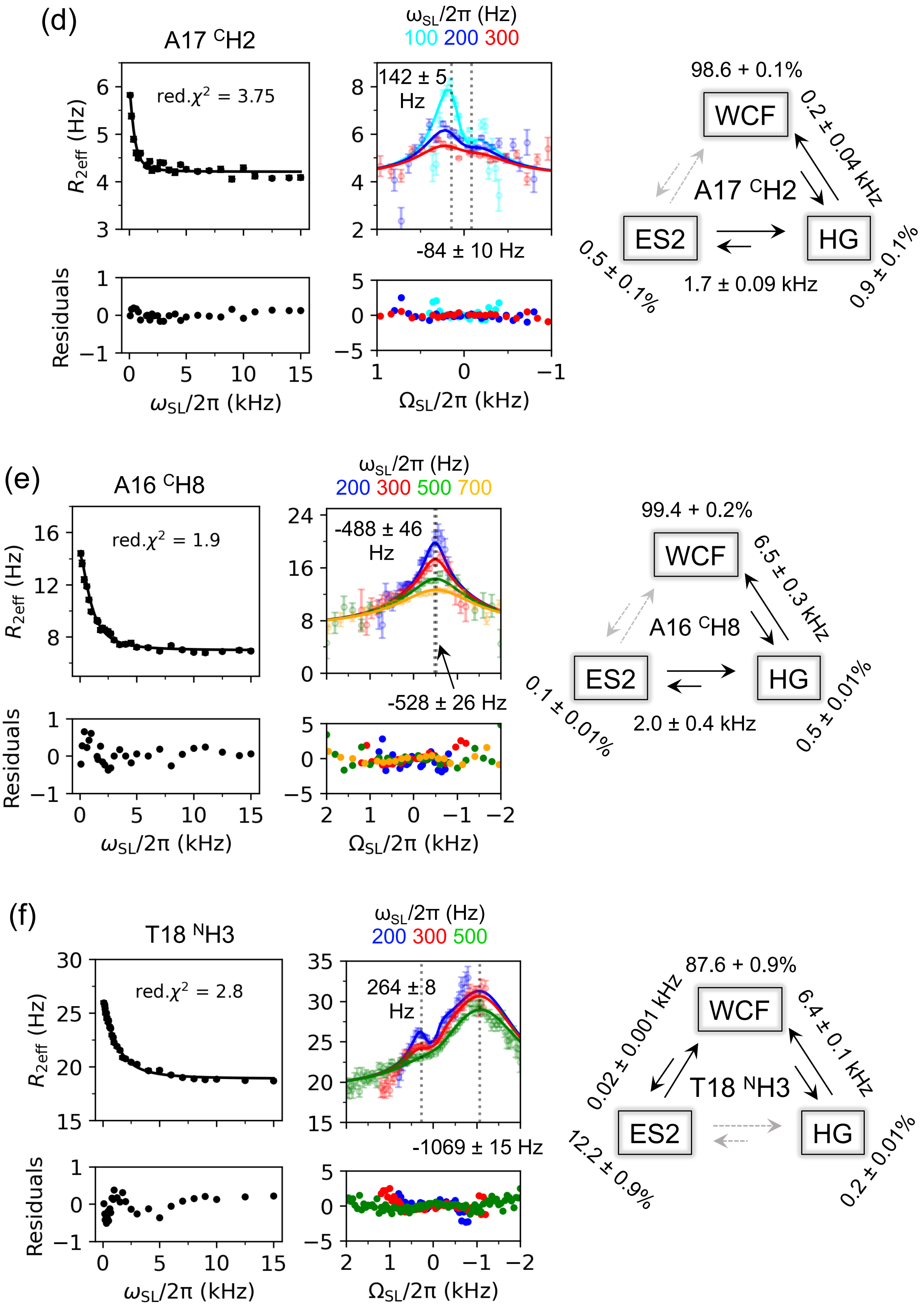
(d)
A17 CH2
red.χ2 = 3.75
R2eff (Hz)
Residuals
ωSL/2π (kHz)
ωSL/2π (Hz)
100 200 300
142 ± 5 Hz
-84 ± 10 Hz
ΩSL/2π (kHz)
98.6 + 0.1%
WCF
0.2 ± 0.04 kHz
A17 CH2
ES2
HG
0.5 ± 0.1%
1.7 ± 0.09 kHz
0.9 ± 0.1%
(e)
A16 CH8
red.χ2 = 1.9
R2eff (Hz)
Residuals
ωSL/2π (kHz)
ωSL/2π (Hz)
200 300 500 700
-488 ± 46 Hz
-528 ± 26 Hz
ΩSL/2π (kHz)
99.4 + 0.2%
WCF
6.5 ± 0.3 kHz
A16 CH8
ES2
HG
0.1 ± 0.01%
2.0 ± 0.4 kHz
0.5 ± 0.01%
(f)
T18 NH3
red.χ2 = 2.8
R2eff (Hz)
Residuals
ωSL/2π (kHz)
ωSL/2π (Hz)
200 300 500
264 ± 8 Hz
-1069 ± 15 Hz
ΩSL/2π (kHz)
87.6 + 0.9%
WCF
0.02 ± 0.001 kHz
6.4 ± 0.1 kHz
T18 NH3
ES2
HG
12.2 ± 0.9%
0.2 ± 0.01%

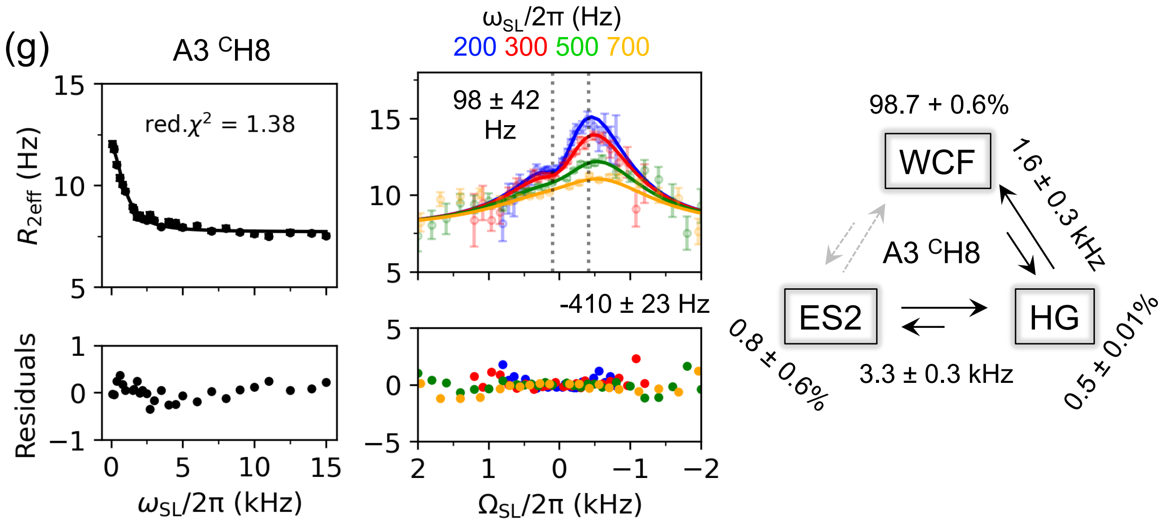


**Figure S3**. **$^1$H $R_{1\rho}$ RD reveals ES2 in WCF – HG dynamics in A$_2$ DNA**. On-resonance (left) and off-resonance (right) $R_{2eff}$ plots for various protons (a) T9 $^N$H3, (b) T8 $^N$H3, (c) A16 $^C$H2, (d) A17 $^C$H2, (e) A16 $^C$H8, and (f) T18 $^N$H3, at 298 K and pH 6.5 showing three-state exchange fits to the data. Solid lines represent the fits associated with the fit parameters listed in Table S1. $\Delta\omega_{HG}$ and $\Delta\omega_{ES2}$ are shown in the off-resonance plots, while exchange rates and populations for each conformer are shown in the schematic representation of the three-state exchange model. Global fits are shown for 1) A16 $^C$H2 and A16 $^C$H8 with shared $k_{ex}$ (WCF – HG) and $p_{HG}$ as well as for 2) T8 $^N$H3 and A17 $^C$H2 with shared $k_{ex}$ (HG – ES2) and $p_{ES2}$. $\Delta\omega_{SL}/2\pi$ used for each off-resonance experiment are color-coded and shown above each plot. Reduced $\chi^2$ values are indicated in the on-resonance plots with the residuals plotted below.

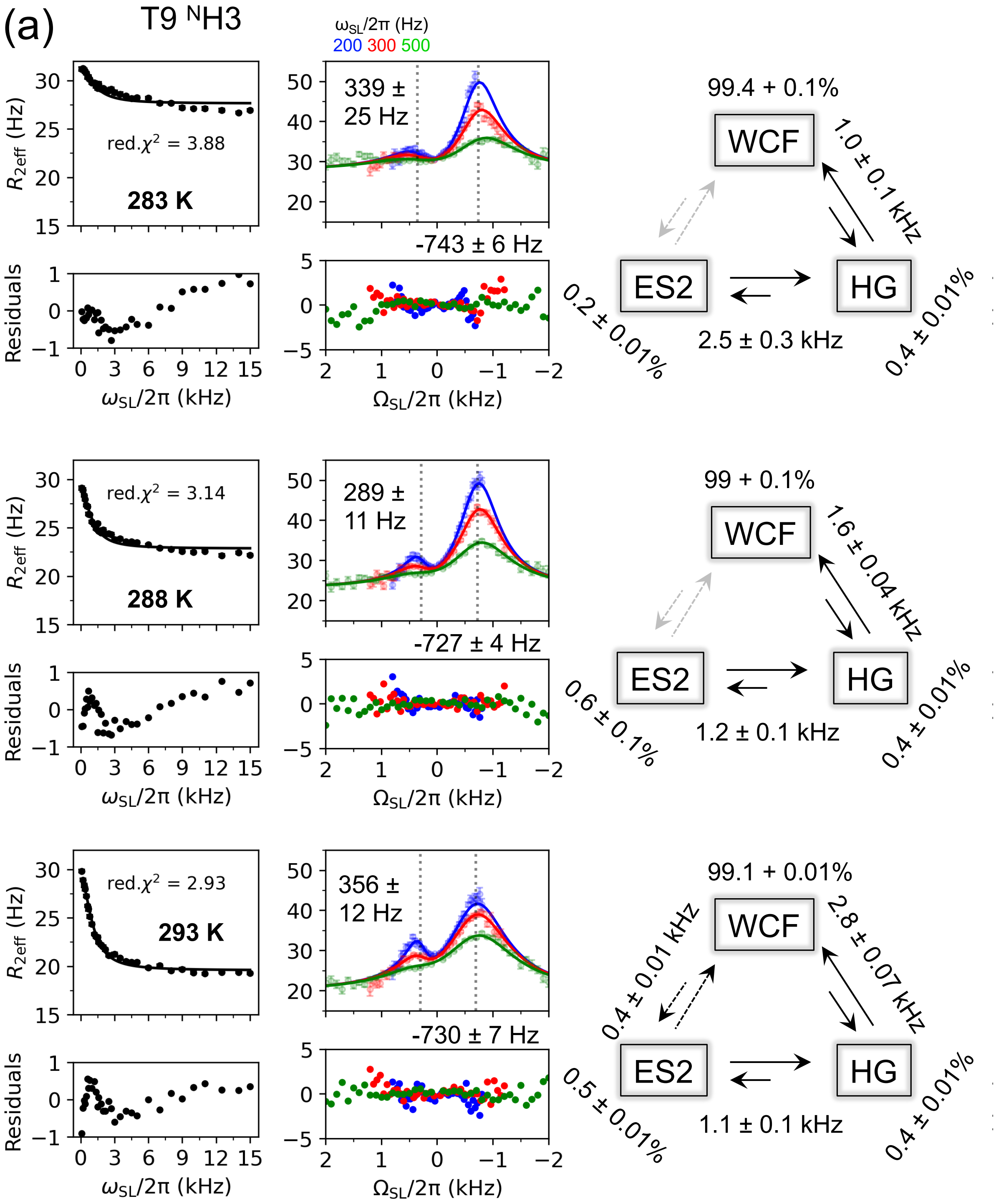
(a)
T9 NH3
ωSL/2π (Hz)
200 300 500
red.χ2 = 3.88
283 K
339 ±
25 Hz
-743 ± 6 Hz
99.4 + 0.1%
WCF
ES2
HG
1.0 ± 0.1 kHz
2.5 ± 0.3 kHz
0.2 ± 0.01%
0.4 ± 0.01%
red.χ2 = 3.14
288 K
289 ±
11 Hz
-727 ± 4 Hz
99 + 0.1%
1.6 ± 0.04 kHz
1.2 ± 0.1 kHz
0.6 ± 0.1%
0.4 ± 0.01%
red.χ2 = 2.93
293 K
356 ±
12 Hz
-730 ± 7 Hz
99.1 + 0.01%
0.4 ± 0.01 kHz
2.8 ± 0.07 kHz
1.1 ± 0.1 kHz
0.5 ± 0.01%
0.4 ± 0.01%
R2eff (Hz)
Residuals
ωSL/2π (kHz)
ΩSL/2π (kHz)

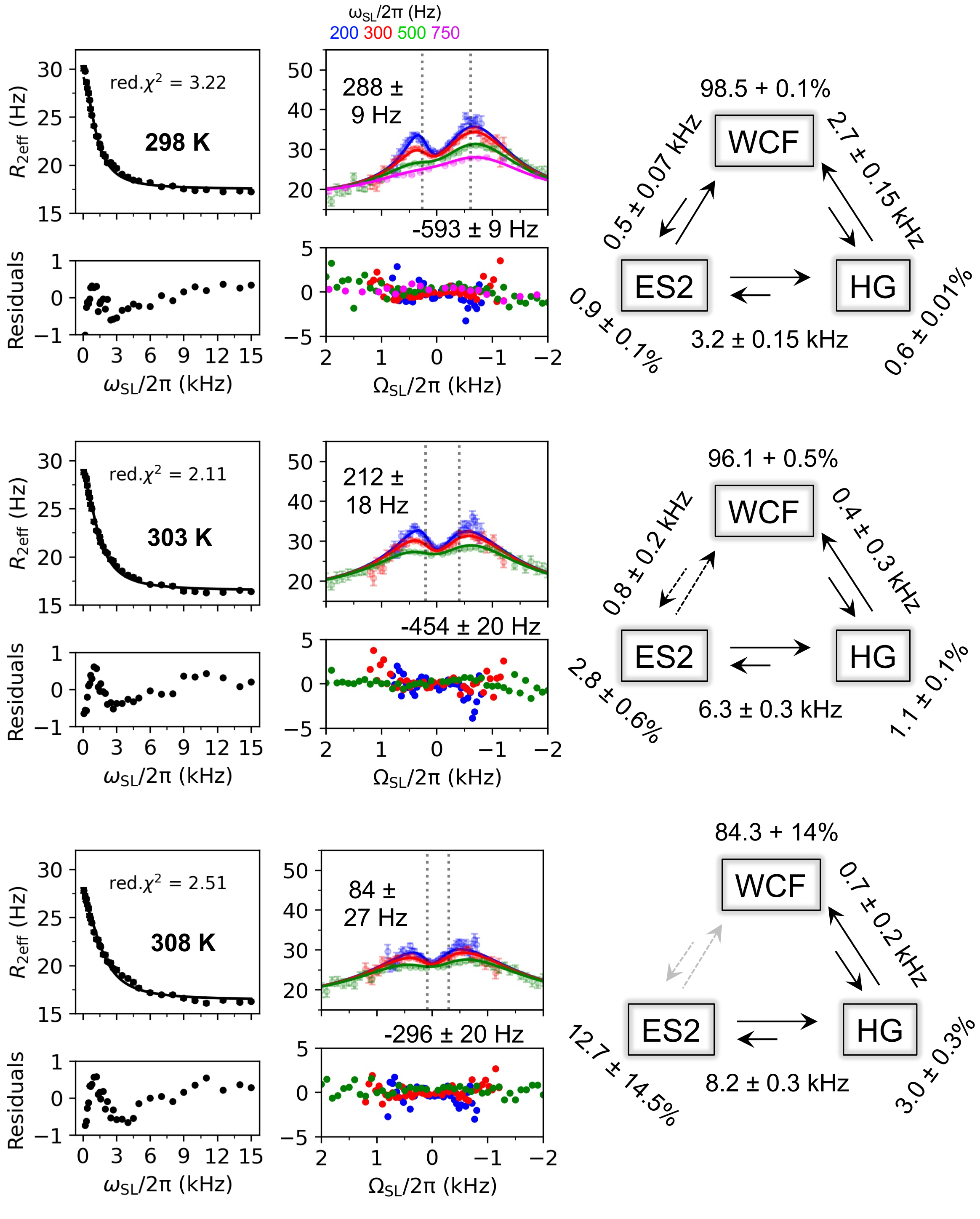

ωSL/2π (Hz)
200 300 500 750
red.χ2 = 3.22
298 K
R2eff (Hz)
Residuals
ωSL/2π (kHz)
288 ±
9 Hz
-593 ± 9 Hz
ΩSL/2π (kHz)
98.5 + 0.1%
WCF
ES2
HG
0.5 ± 0.07 kHz
2.7 ± 0.15 kHz
3.2 ± 0.15 kHz
0.9 ± 0.1%
0.6 ± 0.01%
red.χ2 = 2.11
303 K
212 ±
18 Hz
-454 ± 20 Hz
96.1 + 0.5%
0.8 ± 0.2 kHz
0.4 ± 0.3 kHz
6.3 ± 0.3 kHz
2.8 ± 0.6%
1.1 ± 0.1%
red.χ2 = 2.51
308 K
84 ±
27 Hz
-296 ± 20 Hz
84.3 + 14%
0.7 ± 0.2 kHz
8.2 ± 0.3 kHz
12.7 ± 14.5%
3.0 ± 0.3%

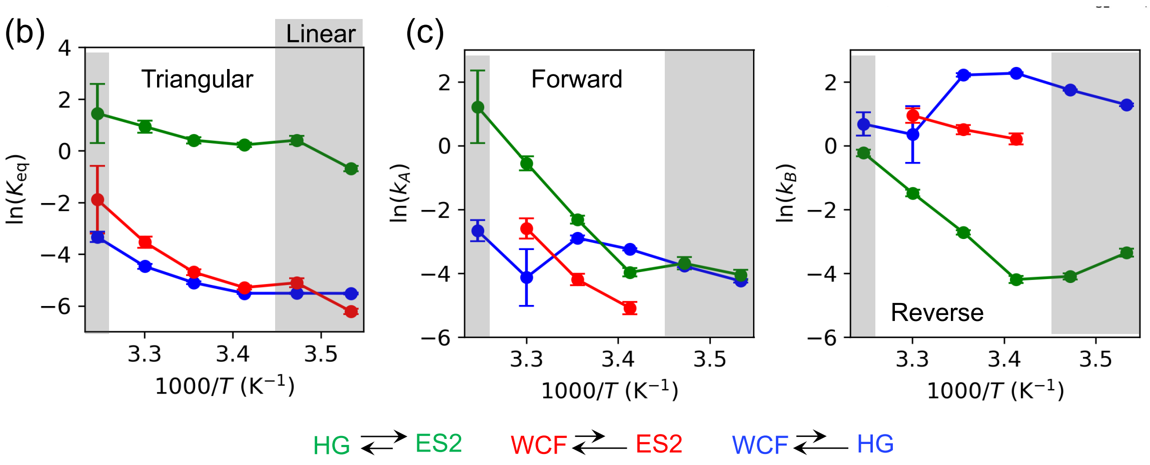


**Figure S4**. (a) Temperature-dependent $^{1}$H $R_{1\rho}$ relaxation dispersion data for T9 $^{N}$H3 in $A_2$ DNA obtained at temperatures ranging from 283 to 308 K in 5 K increments. The solid lines indicate fits using a three-state exchange model, with the exchange parameters and topology depicted (refer to supporting Excel file for model selection information and other fits). The $\omega_{SL}/2\pi$ used for off-resonance experiments at each temperature are color-coded. Reduced $\chi^2$ values and corresponding temperatures are annotated on the on-resonance plots, with the fit residuals displayed below each curve. (b) The van't Hoff plots of ln($K_{eq}$) vs inverse temperature with $K_{eq}$ = $p_{HG}/p_{WCF}$ or $p_{ES2}/p_{WCF}$ or $p_{ES2}/p_{HG}$ for WCF – HG (blue), WCF – ES2 (red) and HG – ES2 (green) transition. (c) Arrhenius plots for each transition, where the forward and reverse rate constants are denoted as $k_A$ and $k_B$ and are calculated from the exchange rates and populations. For the WCF – ES2 transition, forward and reverse rates could only be calculated for temperatures 293, 298 and 303 K because for other temperatures the experiment was not sensitive to detect this transition. This resulted in the linear topology of the three-state exchange model to be favoured for temperatures 283, 288 and 308 K by the model selection criteria (indicated in grey), largely to a too fast exchange process in relation to size of $p_{ES2}$. This multi-topology behaviour of the $^{1}$H $R_{1\rho}$ temperature dependence complicates the estimation of thermodynamics parameters from either the van't Hoff or the Arrhenius plots which cannot be consolidated with a single thermodynamic model.

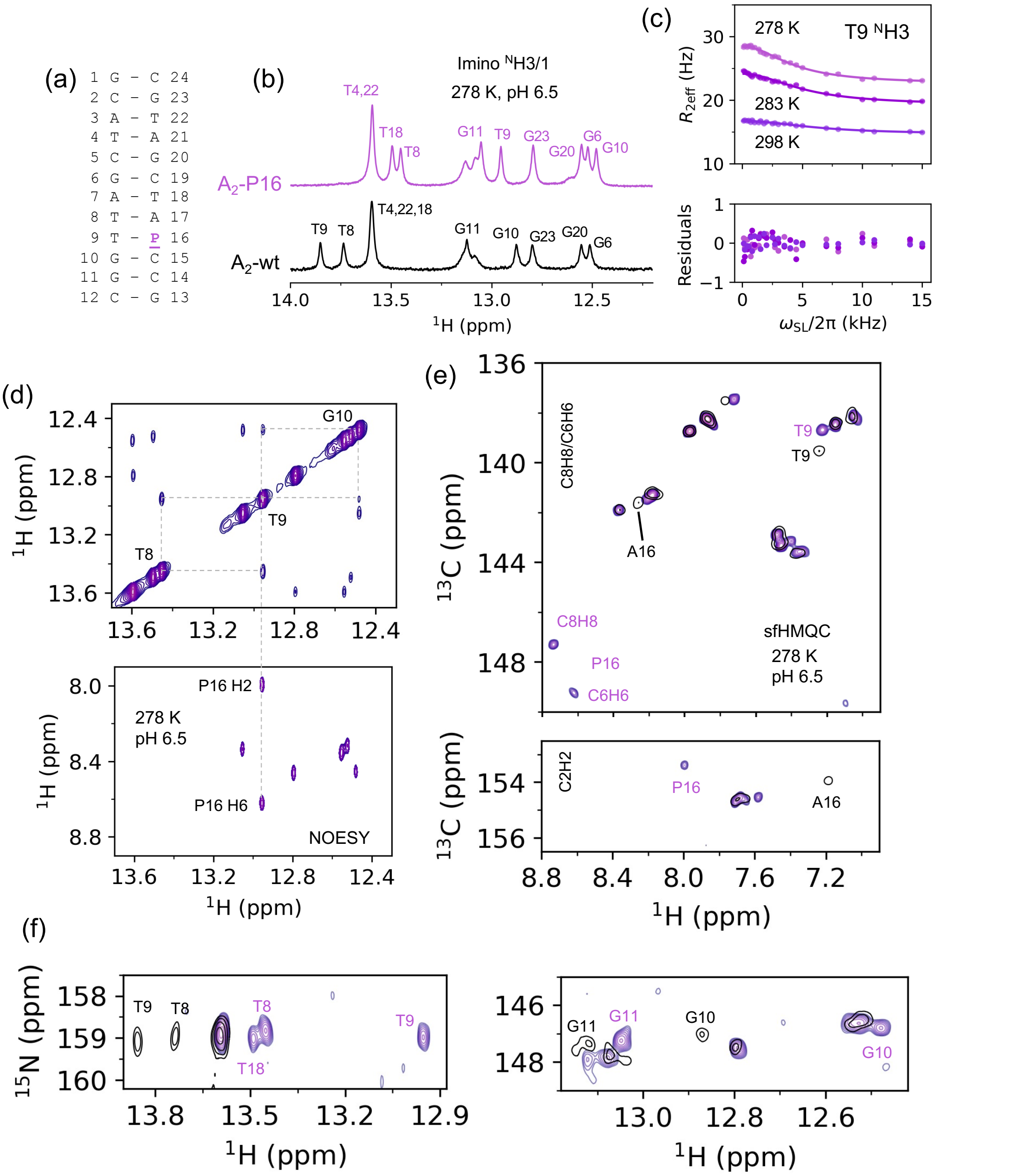


**Figure S5**. **NMR of A2-P16 at 278 K and pH 6.5**. (a) Secondary structure of A2-P16 DNA where the modified base is denoted with bold violet P. (b) 1D $^{1}$H imino spectrum comparing A2-P16 (violet) and A2 wt (black) with resonance assignment denoted. (c) Temperature-dependent $^{1}$H $R_{1\rho}$ on-resonance profiles of T9 $^{N}$H3 in A2-P16 are shown at 278 K, 283 K and 298 K. Solid lines show the fit of two-state reduced exchange model with the residuals plotted below with fit parameters reported in Table S4. It is evident that at 298 K the exchange rate is too fast while at 278 K the exchange regime becomes accessible to study with $^{1}$H $R_{1\rho}$. (d) $^{1}$H–$^{1}$H NOESY spectrum of the imino (top) and aromatic region (bottom) with a mixing time of 180 ms. Cross peaks from T9 $^{N}$H3 to P16 H2 and P16 H6 are shown to support the predominance of the WCF conformation. (e) and (f) Display the SOFAST-HMQC spectra for $^{1}$H–$^{13}$C (C2H2, C6H6 and C8H8) and $^{1}$H–$^{15}$N (imino), respectively, for both A2-wt (black) and A2-P16 (violet) along with the relevant assignments (Table S7).

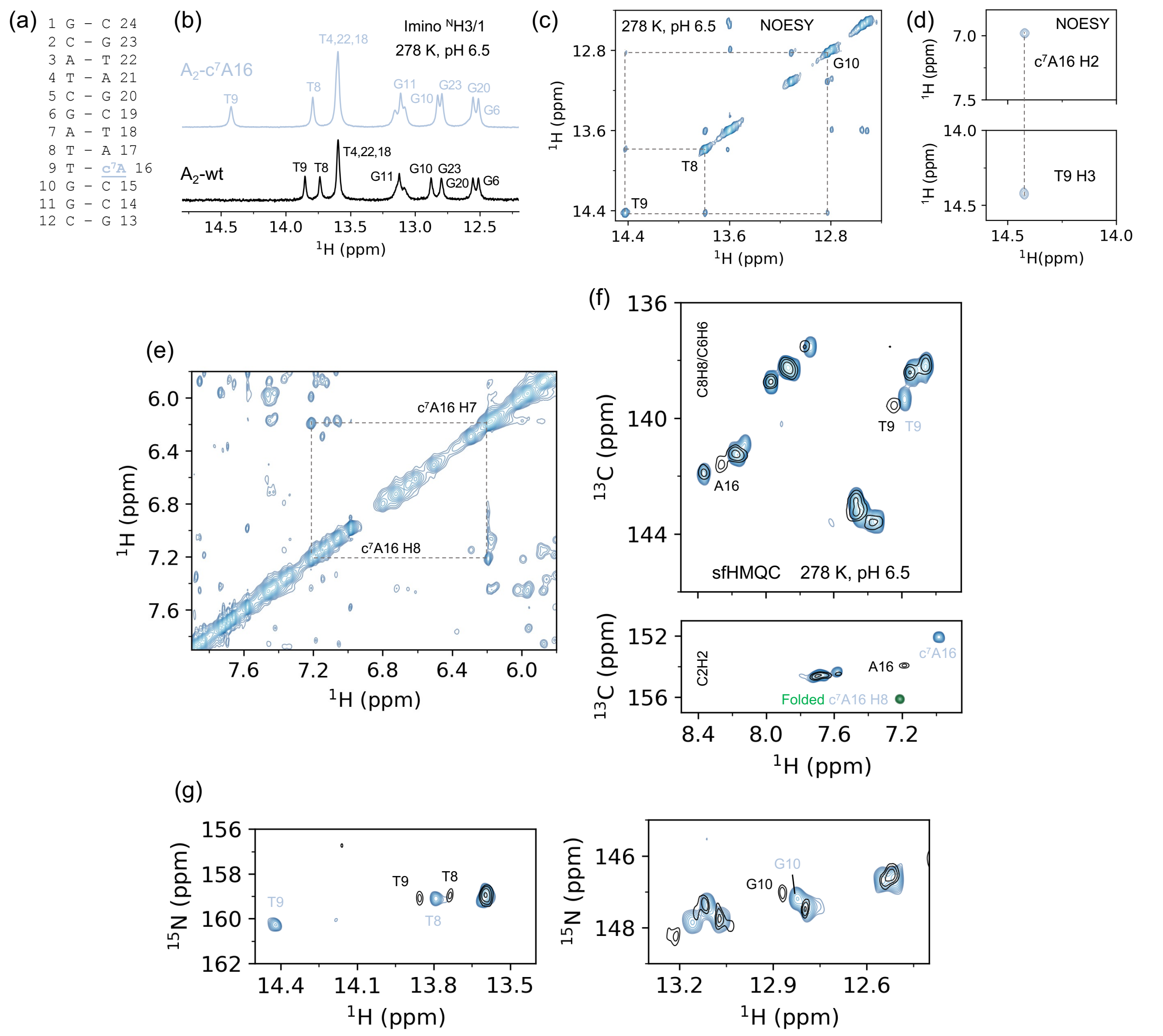


**Figure S6**. **NMR of A2-c⁷A16 at 278 K and pH 6.5**. (a) Secondary structure of A2-c7A16 where the modified nucleotide is denoted in bold blue. (b) 1D $^{1}$H imino spectrum with resonance assignment comparing A2-c$^{7}$A16 (blue) and A2 wt (black). (c) $^{1}$H-$^{1}$H NOESY spectra showing the imino walk for T8, T9 and G10 in grey dashed lines confirm the resonance assignment. (d) A NOESY cross-peak between T9 $^{N}$H3 and c$^{7}$A16 H2 confirms WCF conformation for this base pair. (e) NOESY spectrum shows the expected cross-peaks between c$^{7}$A16 H8 and H7. (f) and (g) Display the SOFAST-HMQC spectra for $^{1}$H–$^{13}$C (C2H2, C6H6 and C8H8) and $^{1}$H–$^{15}$N (imino), respectively, for both A2-wt (black) and A2-c$^{7}$A16 (t blue) with partial assignment for relevant nucleotides consistent with the literature[25]. Resonance assignments are reported in Table S8.

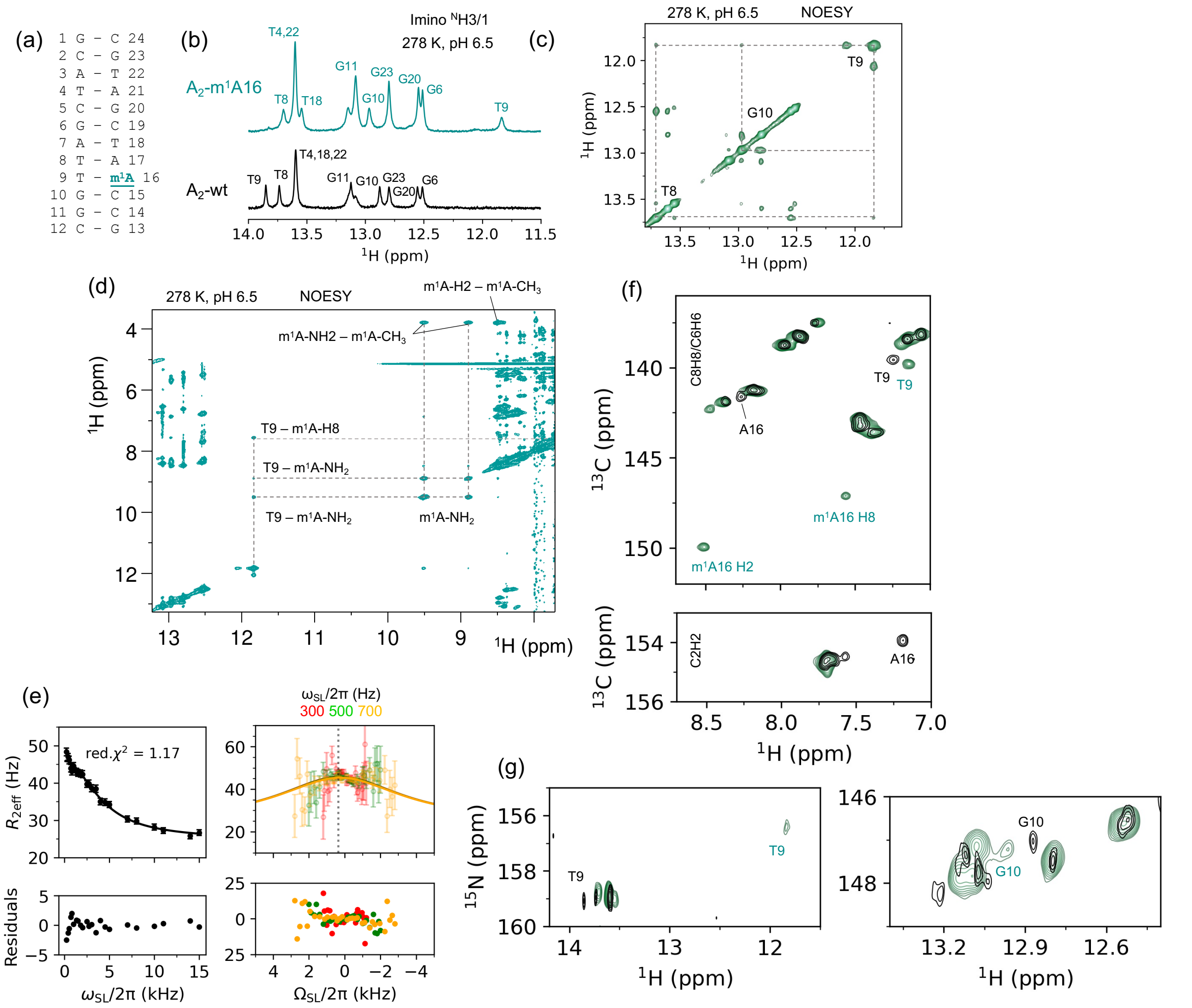

**Figure S7**. **NMR of A2-m$^1$A16 at 278 K and pH 6.5**. (a) Secondary structure of A2 m$^1$A16 where the modified nucleotide is shown in bold green. (b) 1D $^1$H imino spectra and resonance assignment comparing A2-m$^1$A16 (green) and A2-wt (black) (c) $^1$H–$^1$H NOESY walk connecting imino protons from T8, T9 and G10 confirms the resonance assignment. (d) NOESY cross-peak between T9 $^N$H3 and m$^1$A16 H8, confirming the HG conformation for this base pair. The expected cross-peaks between the m$^1$A -NH2 and -CH3 groups are also observed. (e) The $^1$H $R_{1\rho}$ experiment, performed on T9 $^N$H3 at 278 K for A2-m$^1$A16, shows that while the on-resonance profile displays significant relaxation dispersion, the off-resonance profile fails to capture relevant exchange parameters due to fast dynamics (Table S4 and supporting Excel file). The solid line represents a two-state exchange model fit, with the reduced $\chi^2$ value displayed on the on-resonance profile (right), accompanied by the residuals plotted below each curve. The dashed line on the off-resonance plot (left) represents the $\Delta\omega_{ES}$ from the two-state exchange model (f, g) Displays the SOFAST-HMQC spectra for $^1$H–$^{15}$N (imino) and $^1$H–$^{13}$C (C2H2, C6H6 and C8H8) respectively, for both A2-wt (black) and A2-m$^1$A (green). Relevant resonance assignments (refer to Table S9) are denoted, consistent with previous studies[10,26].

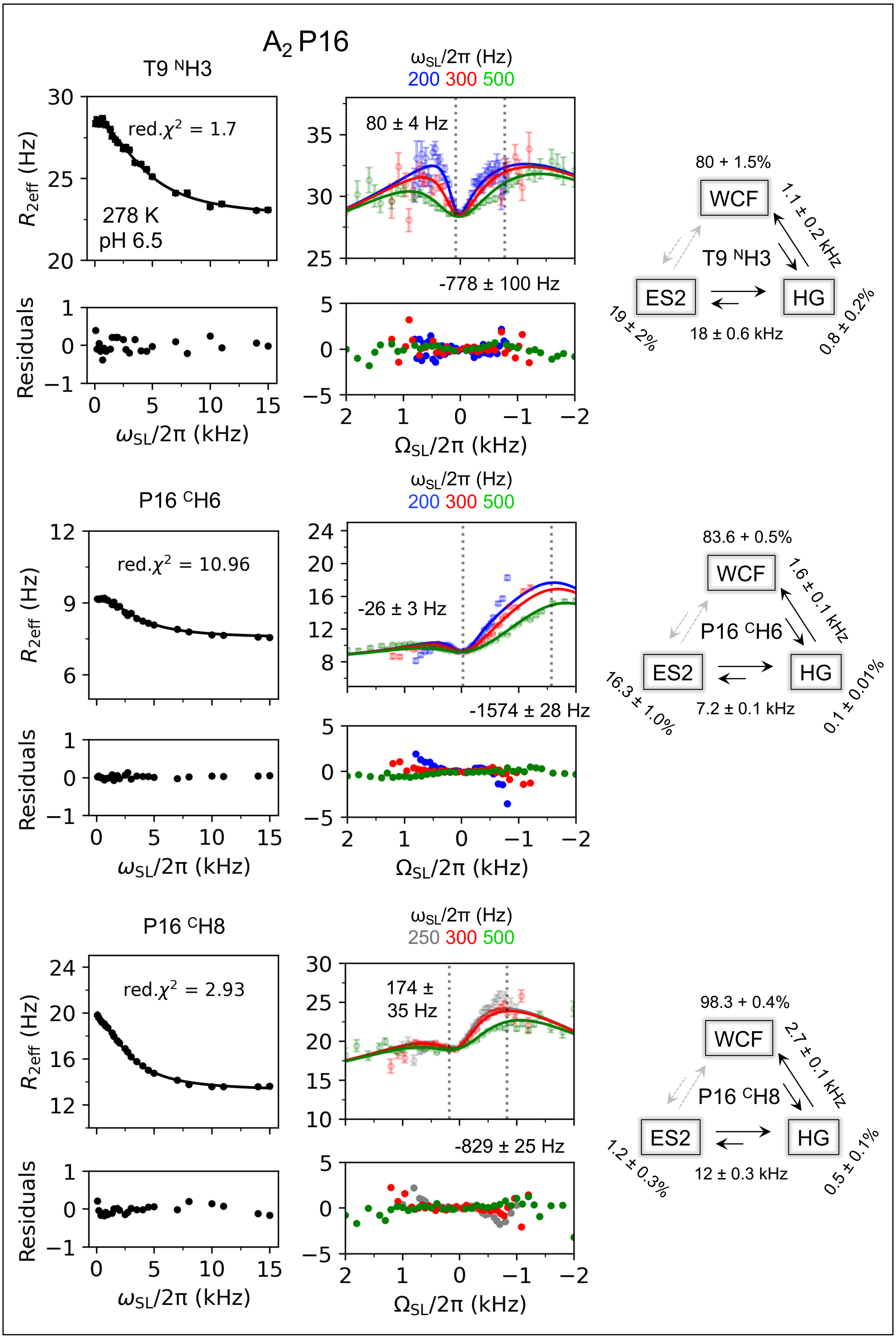
A2 P16
T9 NH3
red.χ2 = 1.7
278 K
pH 6.5
R2eff (Hz)
Residuals
ωSL/2π (kHz)
ωSL/2π (Hz)
200 300 500
80 ± 4 Hz
-778 ± 100 Hz
ΩSL/2π (kHz)
80 + 1.5%
WCF
1.1 ± 0.2 kHz
T9 NH3
ES2
HG
19 ± 2%
18 ± 0.6 kHz
0.8 ± 0.2%
P16 CH6
red.χ2 = 10.96
-26 ± 3 Hz
-1574 ± 28 Hz
83.6 + 0.5%
1.6 ± 0.1 kHz
16.3 ± 1.0%
7.2 ± 0.1 kHz
0.1 ± 0.01%
P16 CH8
red.χ2 = 2.93
250 300 500
174 ± 35 Hz
-829 ± 25 Hz
98.3 + 0.4%
2.7 ± 0.1 kHz
1.2 ± 0.3%
12 ± 0.3 kHz
0.5 ± 0.1%

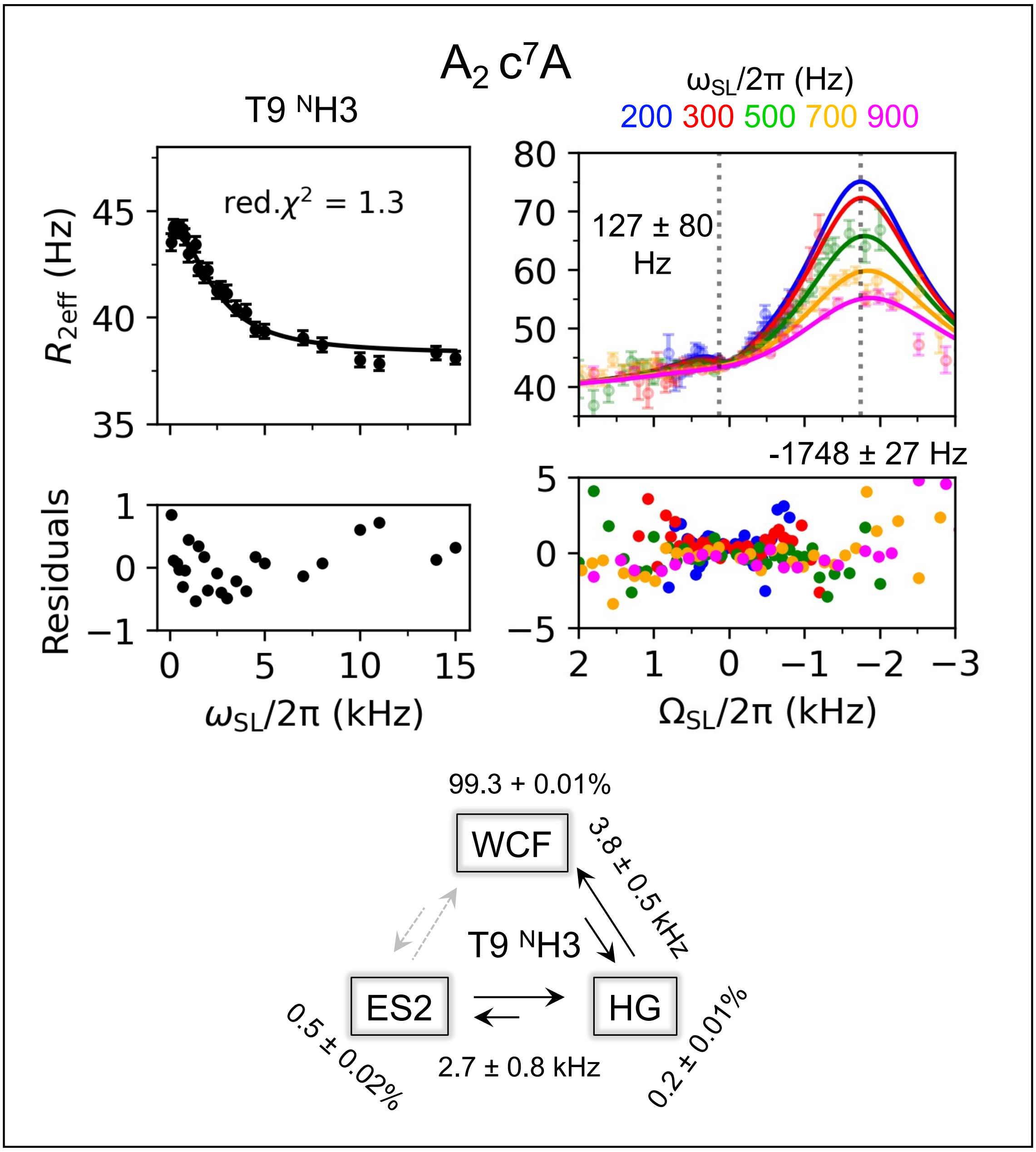


**Figure S8**. **Effect of modified adenine in the A16-T9 base pair on the WCF–HG–ES2 dynamics.** $^{1}$H $R_{1\rho}$ plots at 278 K and pH 6.5 for T9 $^{N}$H3, P16 $^{C}$H6, P16 $^{C}$H8 in A$_2$ P16, and T9 $^{N}$H3 in A$_2$-c$^7$A16. The solid lines indicate fits using a three-state linear exchange model (see supporting Excel file for model selection information). The reduced $\chi^2$ values are shown on the on-resonance plots, while the fit residuals displayed below each curve. The values of $\Delta\omega_{HG}$ and $\Delta\omega_{ES2}$ are highlighted on the off-resonance plots with dotted black lines. Due to high $k_{ex}$ (HG – ES2), $p_{ES2}$, $k_{ex}$ (HG – ES2), and $\Delta\omega_{ES2}$ become correlated and have larger errors than displayed (see MC plots). The exchange rates and populations are summarized in the schematic representation of the three-state exchange model. These results indicate the presence of the ES2 across both modified DNA constructs, with the detection of the HG-like state in the WCF-trapped A$_2$-c$^7$A16 duplex.

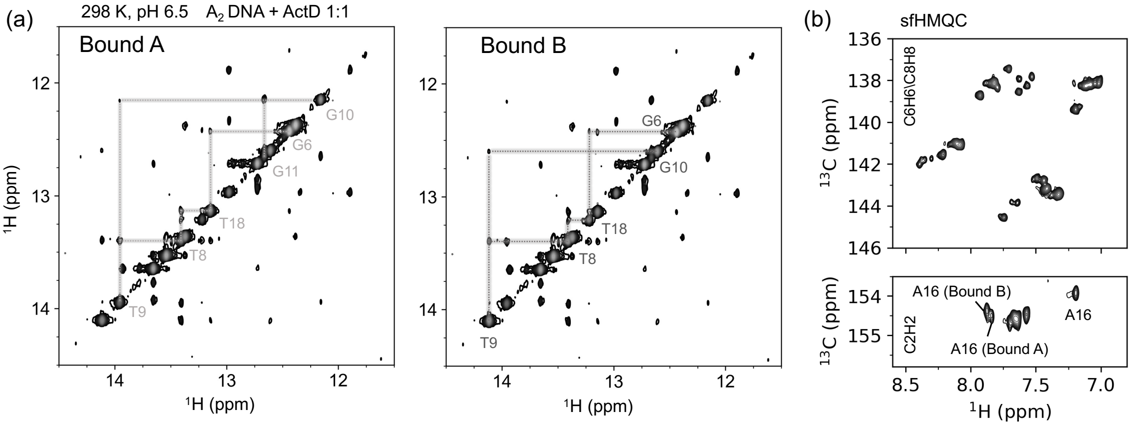


**Figure S9**. **Characterization of Actinomycin D (ActD) bound $A_2$-DNA**. (a) Imino proton NOESY walk for the two ActD-bound fractions (Bound A and Bound B), supporting the assignment in Figure 5a. Cross-peaks confirm sequential connectivity, aiding in the identification of key resonance in the bound states. (b) $^1H - ^{13}C$ SOFAST-HMQC[2] spectrum depicting the A16 $^{C}H2$ chemical shift for both Bound A and Bound B conformations. Although spectral complexity and overlap limited full resonance assignment, the observed peaks provide partial confirmation of A16 involvement in both binding conformations.

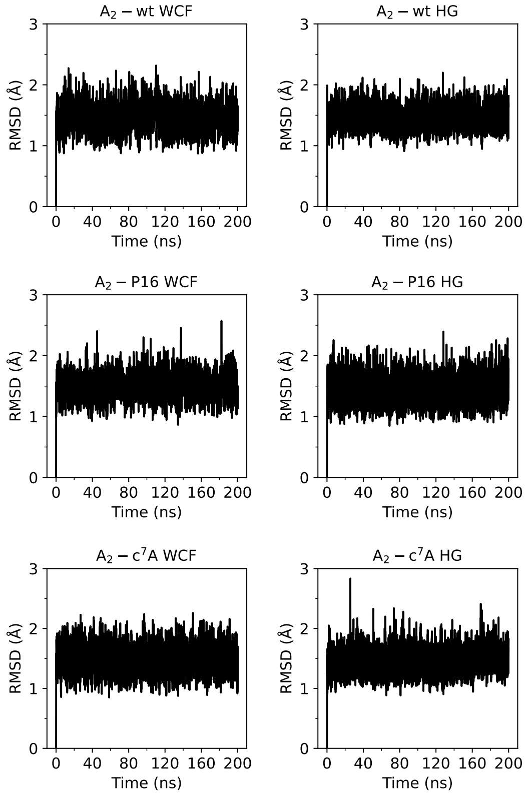


**Figure S10**. **Backbone RMSD of $A_2$-wt, $A_2$-P16, and $A_2$-c$^7$A16 in WCF and HG conformations.**
Root-mean-square deviations (RMSD) of the DNA backbone atoms over 200 ns molecular dynamics simulations for $A_2$-wt, $A_2$-P16, and $A_2$-c$^7$A16 in both Watson–Crick (WCF) and Hoogsteen (HG) conformations. The trajectories show overall structural stability of the DNA duplexes within the accuracy and limitations of the OL15 force field, supporting the reliability of subsequent structural and chemical shift analyses.

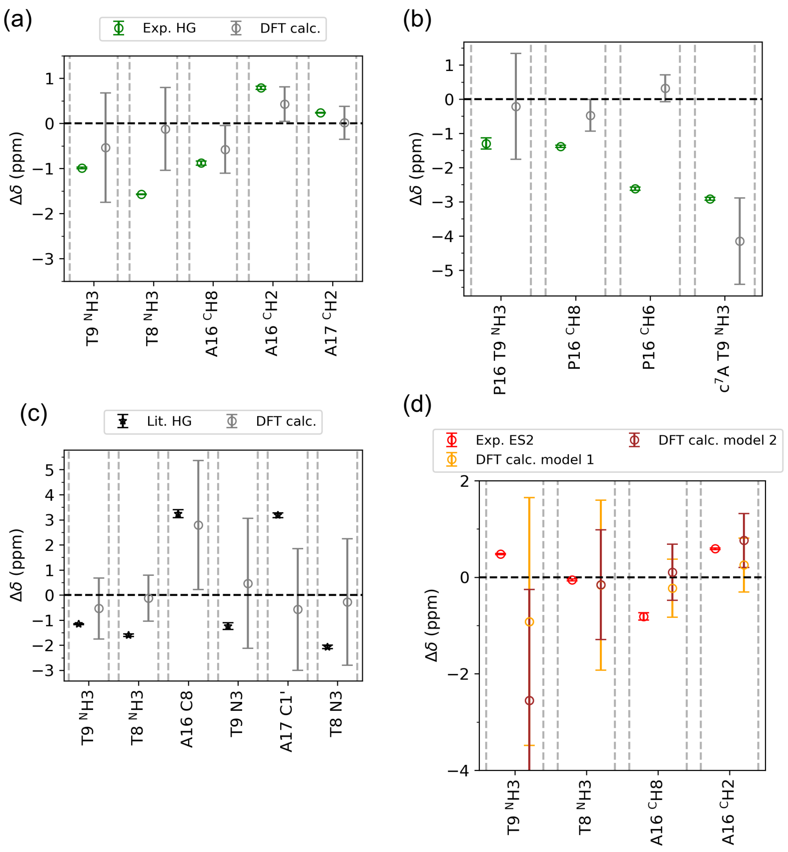


**Figure S11. Benchmarking the chemical shift calculations.** (a) Relative chemical shifts (Δδ, ppm) between HG and WCF conformations obtained from experiments (green) and DFT prediction using AFNMR[20] (grey), based on 50 randomly selected frames from MD simulations. Exchangeable protons (T9 $^{N}$H3, and T8 $^{N}$H3) exhibit higher variability in the calculated Δδ values, whereas non-exchangeable proton (A16 $^{C}$H8, A16 $^{C}$H2, and A17 $^{C}$H2) display more consistent predictions. (b) Δδ values for selected protons—T9 $^{N}$H3, P16 $^{C}$H8, and P16 $^{C}$H6 in $A_2$-P16, and T9 $^{N}$H3 in $A_2$-c$^{7}$A—show good agreement with experimental HG shifts, except for P16 $^{C}$H6, for which the predicted value deviates from the observed trend. (c) Comparison of calculated Δδ values from the same 50 MD-derived frames as in (a) with literature-reported experimental chemical shifts for $^{1}$H, $^{13}$C, and $^{15}$N nuclei[25,27,28]. (d) Experimental Δδ values for ES2 (red) are compared with those calculated for Model 1 (orange) and Model 2 (brown) derived from metadynamics-based clustering. Non-exchangeable protons (A16 $^{C}$H8 and $^{C}$H2) show relatively low variability and better alignment with experimental ES2 shifts, while exchangeable protons (T9 $^{N}$H3 and T8 $^{N}$H3) are less consistent. Although neither model fully captures all features of ES2, additional evidence presented in the main text supports Model 1 as the more likely structural representation.

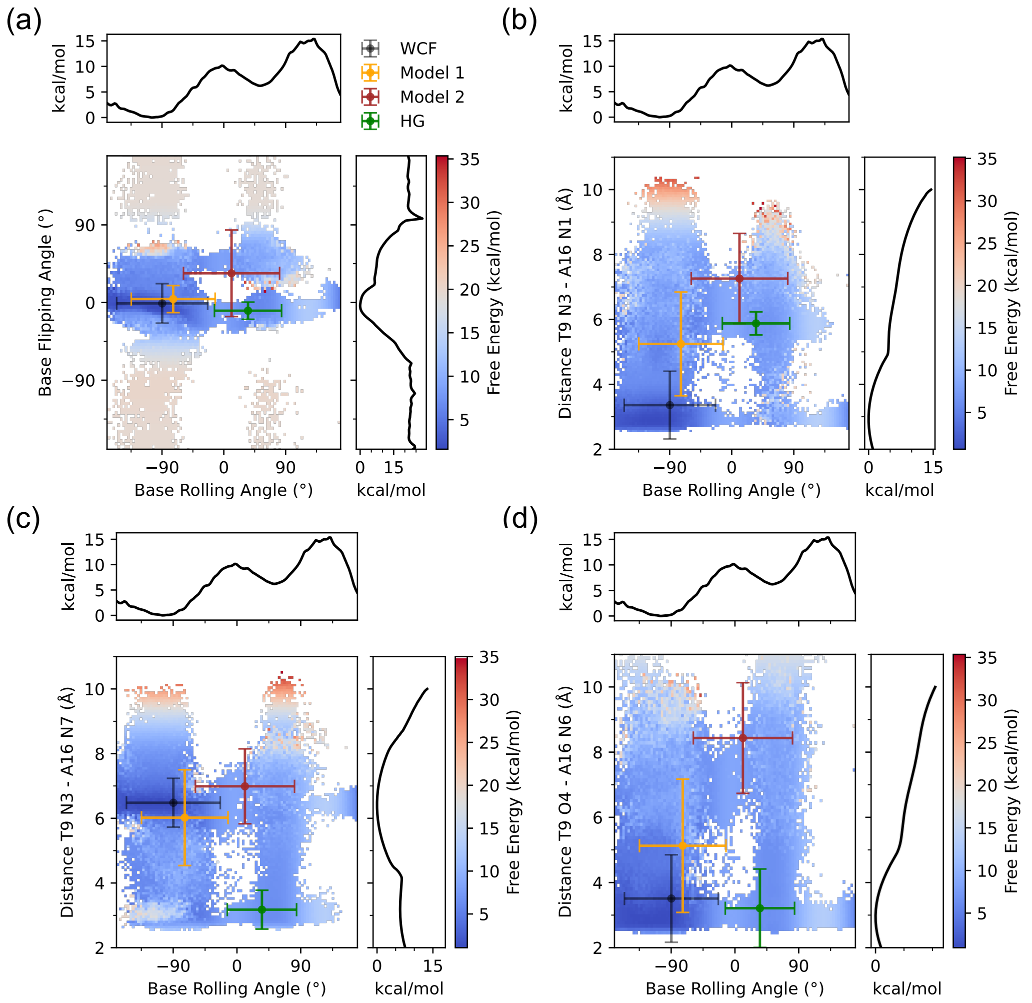


**Figure S12. 2D Free energy surfaces from metadynamics simulations.** Energy-weighted two-dimensional free energy surfaces (2D FES) were constructed from metadynamics simulations using combinations of collective variables (CVs): base rolling angle versus (a) base flipping angle, (b) distance between T9 N3 and A16 N1, (c) distance between T9 N3 and A16 N7, and (d) distance between T9 O4 and A16 N6. Corresponding one-dimensional FES for each CV are shown along the top and right axes of each panel. Clustering based on chemical shift calculations identified four representative structural states: Watson–Crick-Franklin (WCF, black), Hoogsteen (HG, green), and two additional intermediates—Model 1 (orange) and Model 2 (brown). When mapped onto the 2D FES, WCF and HG conformations occupy well-defined minima, consistent with expected stable states. Model 1 consistently localizes to a distinct low-energy basin, suggesting it represents an intermediate along the WCF–HG exchange pathway. In contrast, Model 2 resides in a higher-energy region, indicating a less favourable state compared to Model 1. These observations support Model 1 as the more likely structural candidate for the ES2 intermediate detected in experimental studies.

**Table S1**. Three-state exchange parameters for various protons from the T–A base pair in $A_2$ DNA. The merged cells represent parameters shared in global fiting. The definition of each parameter is given in the main text.

<table>
<tr><th></th><th>T9 $^{N}$H3</th><th>A16 $^{C}$H8</th><th>A16 $^{C}$H2</th><th>T8 $^{N}$H3</th><th>A17 $^{C}$H2</th><th>T18 $^{N}$H3</th><th>A3 $^{C}$H8</th></tr>
<tr><td>$k_{ex}$ ($s^{-1}$)<br>(WC - HG)</td><td>2756 ± 152</td><td colspan="2">6568 ± 303</td><td>1915 ± 22</td><td>276 ± 43</td><td>6378 ± 128</td><td>1636 ± 276</td></tr>
<tr><td>$p_{HG}$ (%)</td><td>0.60 ± 0.01</td><td colspan="2">0.50 ± 0.01</td><td>0.40 ± 0.01</td><td>0.90 ± 0.1</td><td>0.20 ± 0.01</td><td>0.50 ± 0.01</td></tr>
<tr><td>$\Delta\omega_{HG}$ (Hz)</td><td>−593 ± 9</td><td>−528 ± 26</td><td>475 ± 22</td><td>−943 ± 3</td><td>142 ± 5</td><td>−1069 ± 16</td><td>−410 ± 23</td></tr>
<tr><td>$k_{ex}$ ($s^{-1}$)<br>(WC - ES2)</td><td>501 ± 72</td><td>--</td><td>--</td><td>--</td><td>--</td><td>24 ± 0.65</td><td>--</td></tr>
<tr><td>$k_{ex}$ ($s^{-1}$)<br>(HG - ES2)</td><td>3277 ± 155</td><td>2034 ± 431</td><td>1360 ± 242</td><td colspan="2">1723 ± 95</td><td>--</td><td>3339 ± 265</td></tr>
<tr><td>$p_{ES2}$ (%)</td><td>0.90 ± 0.1</td><td>0.10 ± 0.01</td><td>0.30 ± 0.01</td><td colspan="2">0.50 ± 0.10</td><td>12.20 ± 0.90</td><td>0.80 ± 0.6</td></tr>
<tr><td>$\Delta\omega_{ES2}$ (Hz)</td><td>288 ± 9</td><td>−488 ± 46</td><td>354 ± 11</td><td>−33 ± 11</td><td>−84 ± 10</td><td>264 ± 8</td><td>98 ± 42</td></tr>
<tr><td>$R_1$ ($s^{-1}$)</td><td>3.50 ± 0.01</td><td>3.11 ± 0.02</td><td>1.71 ± 0.01</td><td>3.57 ± 0.01</td><td>1.70 ± 0.01</td><td>5.18 ± 0.01</td><td>3.36 ± 0.01</td></tr>
<tr><td>$R_2$ ($s^{-1}$)</td><td>17.10 ± 0.06</td><td>6.96 ± 0.04</td><td>3.91 ± 0.03</td><td>17.3 ± 0.01</td><td>4.21 ± 0.01</td><td>19.00 ± 0.04</td><td>7.71 ± 0.03</td></tr>
<tr><td>Reduced $\chi^2$</td><td>3.24</td><td colspan="2">1.90</td><td colspan="2">3.75</td><td>2.80</td><td>1.38</td></tr>
</table>

**Table S2**. Sequence of unmodified and modified strand of $A_2$ DNA along with the respective yields from solid-phase synthesis.

| DNA oligonucleotide | 5'-sequence-3' | Yield (nmol) |
|---|---|---|
| $A_2$ fwd wt | GCATCGATTGGC | 1073 |
| $A_2$ rev P16 | GCC**X**ATCGATGC | 275 |
| $A_2$ rev $c^7$A16 | GCC**Y**ATCGATGC | 291 |
| $A_2$ rev $m^1$A16 | GCC**Z**ATCGATGC | 248 |
| $A_2$ rev wt* | GCCAATCGATGC | 1000 |

* Obtained from Integrated DNA Technologies, fwd = Forward strand, rev = Reverse strand
X = 2'-deoxynebularine (P), Y = 2'-deoxy-7-deazaadenosine ($c^7$A), Z = 1-methyl-2'-deoxyadenosine ($m^1$A)

**Table S3**. Sample information for NMR spectroscopy of various DNA used in this study at pH 6.5.

| DNA duplex | Conc. for NMR (mM) | Temperature |
|---|---|---|
| $A_2$-wt | 1.5 | 288 – 308 K |
| $A_2$-P16 | 1.1 | 278 – 298 K |
| $A_2$-$c^7$A16 | 1.2 | 278 K |
| $A_2$-$m^1$A16 | 1.0 | 278 K |

**Table S4**. Fit parameters from temperature-dependent on-resonance $^1$H $R_{1\rho}$ RD experiments of T9 $^N$H3 in $A_2$ P16 fitted with a reduced two-state model[29]. Parameters for T9 $^N$H3 in $A_2$ m$^1$A16 fitted with the two-state exchange model at 278 K is also reported.

| T9 $^N$H3 | Fit parameters | 298 K | 283 K | 278 K |
|---|---|---|---|---|
| $A_2$-P16 | $k_{ex}$ (s$^{-1}$) | 39670 ± 3255 | 32470 ± 1244 | 27084 ± 915 |
| | Φ / 4π$^2$ (Hz$^2$) | 2100 ± 257 | 4040 ± 205 | 4031 ± 172 |
| | $R_2$ (s$^{-1}$) | 14.6 ± 0.1 | 19.2 ± 0.1 | 22.6 ± 0.1 |
| | Reduced $\chi^2$ | 1.5 | 2.9 | 1.3 |
| $A_2$-m$^1$A16 | $k_{ex}$ (s$^{-1}$) | | | 26205 ± 1427 |
| | $p_b$ (%) | | | 13.4 ± 12.2 |
| | $\Delta\omega$ (Hz) | | | 347 ± 231 |
| | $R_1$ (s$^{-1}$) | | | 5.09 ± 0.1 |
| | $R_2$ (s$^{-1}$) | | | 25.0 ± 0.6 |
| | Reduced $\chi^2$ | | | 1.171 |

**Table S5**. Fit parameters from $^1$H $R_{1\rho}$ RD experiments for $A_2$-P16 and $A_2$-c$^7$A at 278 K and pH 6.5.

| | $A_2$-P16 | | | $A_2$-c$^7$A16 |
|---|---|---|---|---|
| | T9 $^N$H3 | P16 $^C$H6 | P16 $^C$H8 | T9 $^N$H3 |
| $k_{ex}$ (s$^{-1}$) (WC - HG) | 1145 ± 266 | 1637 ± 61 | 2727 ± 131 | 3792 ± 501 |
| $p_{HG}$ (%) | 0.8 ± 0.2 | 0.1 ± 0.01 | 0.5 ± 0.01 | 0.2 ± 0.01 |
| $\Delta\omega_{HG}$ (Hz) | −778 ± 100 | −1573 ± 28 | −829 ± 20 | −1748 ± 27 |
| $k_{ex}$ (s$^{-1}$) (WC - ES2) | -- | -- | -- | -- |
| $k_{ex}$ (s$^{-1}$) (HG - ES2) | 18370 ± 612 | 7206 ± 133 | 12528 ± 9 | 2686 ± 893 |
| $p_{ES2}$ (%) | 19.3 ± 2.2 | 16.3 ± 1 | 1.2 ± 0.2 | 0.5 ± 0.02 |
| $\Delta\omega_{ES2}$ (Hz) | 80 ± 4.5 | −27 ± 3 | 174 ± 13 | 127 ± 89 |
| $R_1$ (s$^{-1}$) | 3.2 ± 0.1 | 2.1 ± 0.01 | 5.0 ± 0.01 | 16.2 ± 0.1 |
| $R_2$ (s$^{-1}$) | 22.6 ± 0.1 | 7.5 ± 0.01 | 13.2 ± 0.02 | 38.3 ± 0.2 |
| Reduced $\chi^2$ | 1.7 | 10.9 | 2.9 | 1.3 |

**Table S6**. Spin-lock values ($\omega_{SL}$) and frequency offsets ($\Omega_{SL}$) with respect to the signal of interest used to acquire on- and off-resonance relaxation dispersion data. 25–35 datapoints were collected in the indicated ranges using eight spin-lock durations. The maximum spin-lock duration for each resonance was chosen such as to observe a decay to approximately 1/3 of the initial peak intensity at the lowest on-resonance $\omega_{SL}$. A similar experimental setup was used for modified $A_2$ DNA at 278 K.

<table>
<tr><th></th><th></th><th>ω<sub>SL</sub> (Hz)</th><th>Ω<sub>SL</sub> (Hz)</th></tr>
<tr><td rowspan="2">T8 NH3</td><td>On-res</td><td>100 to 15000</td><td>0</td></tr>
<tr><td>Off-res</td><td>200<br>300<br>500</td><td>± 800<br>± 1200<br>± 2000</td></tr>
<tr><td rowspan="2">T9 NH3</td><td>On-res</td><td>100 to 15000</td><td>0</td></tr>
<tr><td>Off-res</td><td>200<br>250<br>300<br>500<br>750</td><td>± 800<br>± 1000<br>± 1200<br>± 2000<br>± 3000</td></tr>
<tr><td rowspan="2">T18 NH3</td><td>On-res</td><td>100 to 15000</td><td>0</td></tr>
<tr><td>Off-res</td><td>200<br>300<br>500</td><td>± 800<br>± 1200<br>± 2000</td></tr>
<tr><td rowspan="2">A16 CH2</td><td>On-res</td><td>100 to 15000</td><td>0</td></tr>
<tr><td>Off-res</td><td>200<br>300<br>500<br>700</td><td>± 800<br>± 1200<br>± 2000<br>± 2800</td></tr>
<tr><td rowspan="2">A16 CH8</td><td>On-res</td><td>100 to 15000</td><td>0</td></tr>
<tr><td>Off-res</td><td>200<br>300<br>500<br>700</td><td>± 800<br>± 1200<br>± 2000<br>± 2800</td></tr>
<tr><td rowspan="2">A17 CH2</td><td>On-res</td><td>100 to 15000</td><td>0</td></tr>
<tr><td>Off-res</td><td>100<br>200<br>300</td><td>± 400<br>± 800<br>± 1200</td></tr>
<tr><td rowspan="2">P16 CH6</td><td>On-res</td><td>100 to 15000</td><td>0</td></tr>
<tr><td>Off-res</td><td>200<br>300<br>500</td><td>± 800<br>± 1200<br>± 2000</td></tr>
<tr><td rowspan="2">P16 CH8</td><td>On-res</td><td>100 to 15000</td><td>0</td></tr>
<tr><td>Off-res</td><td>200<br>300<br>500</td><td>± 800<br>± 1200<br>± 2000</td></tr>
<tr><td rowspan="2">A2-P16<br>T9 NH3</td><td>On-res</td><td>100 to 15000</td><td>0</td></tr>
<tr><td>Off-res</td><td>200<br>300<br>500</td><td>± 800<br>± 1200<br>± 2000</td></tr>
<tr><td rowspan="2">A2-c7A<br>T9 NH3</td><td>On-res</td><td>100 to 15000</td><td>0</td></tr>
<tr><td>Off-res</td><td>200<br>300<br>500<br>700<br>900</td><td>± 800<br>± 1200<br>± 2000<br>±2800<br>±3600</td></tr>
</table>

**Table S7**. Chemical shift assignment in ppm for $A_2$-P16 at 278 K and pH 6.5

| | H3 | H1 | H2 | H8 | H6 | C2 | C8 | C6 | N3 | N1 |
|---|---|---|---|---|---|---|---|---|---|---|
| G1 | | | | 7.970 | | | 138.733 | | | |
| C2 | | | | | | | | | | |
| A3 | | | | 8.370 | | | 141.910 | | | |
| T4 | 13.596 | | | | | | | | | |
| C5 | | | | | | | | | | |
| G6 | | 12.523 | | | | | | | | |
| A7 | | | 7.655 | | | 154.635 | | | | |
| T8 | 13.452 | | | | | | | | 158.825 | |
| T9 | 12.955 | | | | 7.223 | | | 138.692 | 158.985 | |
| G10 | | 12.479 | | | | | | | | 146.803 |
| G11 | | 13.053 | | 7.721 | | | 137.476 | | | 147.250 |
| C12 | | | | | | | | | | |
| G13 | | | | | | | | | | |
| C14 | | | | | | | | | | |
| C15 | | | | | | | | | | |
| P16 | | | 7.995 | 8.737 | 8.621 | 153.385 | 147.309 | 149.243 | | |
| A17 | | | 7.581 | | | 154.544 | | | | |
| T18 | 13.495 | | | | | | | | 159.014 | |
| C19 | | | | | | | | | | |
| G20 | | 12.554 | | | | | | | | |
| A21 | | | | | | | | | | |
| T22 | 13.592 | | | | | | | | | |
| G23 | | 12.794 | | | | | | | | 147.516 |
| C24 | | | | | | | | | | |

**Table S8**. Chemical shift assignment in ppm for $A_2$-$c^7A16$ at 278 K and pH 6.5

| | H3 | H1 | H2 | H8 | H6 | H7 | C2 | C8 | C6 | N3 | N1 |
|---|---|---|---|---|---|---|---|---|---|---|---|
| G1 | | 13.080 | | | | | | | | | 147.798 |
| C2 | | | | | | | | | | | |
| A3 | | | | 8.366 | | | | 141.927 | | | |
| T4 | 13.598 | | | | | | | | | | |
| C5 | | | | | | | | | | | |
| G6 | | 12.513 | | | | | | | | | 146.628 |
| A7 | | | | | | | | | | | |
| T8 | 13.791 | | | | | | | | | 159.104 | |
| T9 | 14.422 | | | | 7.185 | | | | 139.345 | 160.278 | |
| G10 | | 12.826 | | | | | | | | | 147.199 |
| G11 | | 13.112 | | | | | | | | | 147.381 |
| C12 | | | | | | | | | | | |
| G13 | | | | | | | | | | | |
| C14 | | | | | | | | | | | |
| C15 | | | | | | | | | | | |
| $c^7$A16 | | | 6.983 | 7.213 | | 6.20 | 152.14 | | | | |
| A17 | | | 7.580 | | | | 154.373 | | | | |
| T18 | 13.606 | | | | | | | | | 159.082 | |
| C19 | | | | | | | | | | | |
| G20 | | 12.553 | | | | | | | | | 146.676 |
| A21 | | | | | | | | | | | |
| T22 | 13.595 | | | | | | | | | | |
| G23 | | 12.794 | | | | | | | | | 147.447 |
| C24 | | | | | | | | | | | |

**Table S9**. Chemical shift assignment in ppm for $A_2$-m[1]A16 at 278 K and pH 6.5

| | H3 | H1 | H2 | H8 | H6 | C2 | C8 | C6 | N3 | N1 | $NH_2$ (1/2) | $CH_3$ |
|---|---|---|---|---|---|---|---|---|---|---|---|---|
| G1 | | 13.103 | | | | | | | | | | |
| C2 | | | | | | | | | | | | |
| A3 | | | | 8.366 | | | 141.927 | | | | | |
| T4 | 13.608 | | | | | | | | | | | |
| C5 | | | | | | | | | | | | |
| G6 | | 12.510 | | | | | | | | 146.574 | | |
| A7 | | | | | | | | | | | | |
| T8 | 13.707 | | | | | | | | 158.765 | | | |
| T9 | 11.834 | | | | 7.144 | | | 139.786 | 156.402 | | | |
| G10 | | 12.974 | | | | | | | | 147.236 | | |
| G11 | | 13.082 | | | | | | | | 147.178 | | |
| C12 | | | | | | | | | | | | |
| G13 | | 13.154 | | | | | | | | 147.893 | | |
| C14 | | | | | | | | | | | | |
| C15 | | | | | | | | | | | | |
| m[1]A16 | | | 8.496 | 7.558 | | 149.946 | 147.111 | | | | 9.507/8.896 | 3.785 |
| A17 | | | 7.683 | | | | | | | | | |
| T18 | 13.551 | | | | | | | | 159.020 | | | |
| C19 | | | | | | | | | | | | |
| G20 | | 12.550 | | | | | | | | 146.631 | | |
| A21 | | | | | | | | | | | | |
| T22 | 13.602 | | | | | | | | | | | |
| G23 | | 12.802 | | | | | | | | 147.477 | | |
| C24 | | | | | | | | | | | | |

**Table S10**. Chemical shift assignment in ppm for $A_2$ DNA + ActD in a 1:1 mixture at 298 K and pH 6.5. The unbound fraction is marked in blue while the other two bound fractions are marked in red and green

| | H3 | H1 | H2 | C2 |
|---|---|---|---|---|
| T9 | 13.68 / 13.98 / 14.14 | | | |
| T8 | 13.56 / 13.43 / 13.43 | | | |
| T18 | 13.39 / 13.17 / 13.24 | | | |
| G11 | | 13.00 / 12.69 | | |
| G10 | | 12.75 / 12.18 / 12.63 | | |
| G6 | | 12.41 / 12.46 / 12.46 | | |
| A16 | | | 7.24 / 7.89 / 7.92 | 154.06 / 154.64 / 154.51 |

**Table S11**. Average values and the standard deviation of the collective variables for both WCF and HG conformation from unbiased 100ns MD simulation. The WCF – HG transition was probed for A16 in $A_2$ DNA

| Collective Variables | WCF | HG |
|---|---|---|
| $\chi$ dihedral (radian) | −1.77 ± 0.25 | 0.91 ± 0.19 |
| Base Flipping (radian) | −0.06 ± 0.08 | −0.16 ± 0.05 |
| T9 N3 – A16 N7 (Å) | 6.4 ± 0.14 | 3.0 ± 0.31 |
| T9 O4 – A16 N6 (Å) | 3.0 ± 0.2 | 3.0 ± 0.30 |
| T9 N3 – A16 N1 (Å) | 3.0 ± 0.1 | 5.9 ± 0.2 |
| Centre of Mass C15 – A17 (Å) | 7.8 ± 0.4 | 7.8 ± 0.4 |
